# A clinical-stage oncology compound selectively targets drug-resistant cancers

**DOI:** 10.1101/2025.11.26.690878

**Authors:** Kaitlin Long, Debanjan Bhattacharjee, Samuel H. Newman-Stonebraker, Simon Suhr, Brandon Q. Mercado, Emily Scheib, Anthony Tighe, Luciano Romero, Sarah L. Thompson, Erin L. Sausville, Kristen M. John, Linda Julian, Sanat Mishra, Olaf Klingbeil, Punya Gupta, Uditi Bhatt, Allen C. Gao, Sara Ricardo, Christopher R. Vakoc, Beat C. Bornhauser, Steven M. Corsello, Stephen S. Taylor, Jue Chen, Patrick L. Holland, Jason M. Sheltzer

## Abstract

Re-evaluating existing clinical compounds can uncover previously unrecognized mechanisms that reshape a drug’s therapeutic potential. The small molecule Procaspase-Activating Compound 1 (PAC-1) entered oncology testing as a proposed activator of caspase-driven apoptosis. Here, we show that PAC-1-driven cytotoxicity occurs in the absence of executioner caspase expression, demonstrating that its anti-cancer activity occurs via an alternative mechanism. We provide genetic, biochemical, and biophysical evidence demonstrating that PAC-1 functions as a highly selective iron chelator that eliminates cancer cells by disrupting iron homeostasis. Unexpectedly, we discovered that expression of the key chemotherapy-resistance pump MDR1 confers marked hypersensitivity to PAC-1 treatment. While PAC-1 is only weakly effluxed by MDR1 under basal conditions, this process is potentiated when PAC-1 is bound to iron. Consequently, PAC-1 induces progressive iron depletion and selective cytotoxicity in otherwise drug-resistant MDR1-expressing cancer cells. Together, these findings redefine PAC-1’s mechanism-of-action and establish a framework for exploiting multidrug resistance as a therapeutic vulnerability through targeted iron starvation.

## Introduction

Repurposing existing drugs for new clinical indications holds significant promise for precision oncology^1^. Careful genetic and biochemical characterization efforts have revealed previously unknown mechanisms-of-action (MOAs) for existing therapies, which can allow the identification of new biomarkers for patient selection^2–7^. Compounds that have already been demonstrated to be safe in humans can be rapidly tested against new diseases without the investment of hundreds of millions of dollars and years of development typically required for a new therapeutic entity^8^. Alternately, these compounds can serve as advanced starting points for further development based on a novel MOA.

Patients with advanced cancers are frequently treated with cytotoxic chemotherapy, including paclitaxel, doxorubicin, and other agents. However, these compounds are typically able to stop disease progression for only a short period of time as the tumors evolve to become drug-resistant^9^. One common mechanism driving chemotherapy resistance is the upregulation of the drug efflux pump MDR1 (Multidrug Resistance 1, also called ABCB1 or P-glycoprotein)^10–12^. MDR1 encodes a membrane transporter that expels a diverse range of xenobiotics from mammalian cells, including nearly all drugs approved for advanced cancers. Multiple factors contribute to MDR1 upregulation in cancer, including the activation of stress-responsive transcription factors that drive MDR1 expression and the acquisition of intrachromosomal fusion events that place MDR1 under the control of highly-active promoters^12–14^. MDR1 expression severely limits subsequent treatment options, creating an urgent need for new anti-cancer strategies^15^.

Apoptosis, or programmed cell death, is a critical biological pathway for eliminating damaged or malignant cells^16^. The cysteine protease caspase-3 (CASP3) serves as the primary executioner of apoptosis, cleaving hundreds of proteins to induce cell death^17,18^. Caspase-3 normally exists as an inactive zymogen, until it undergoes proteolytic activation stimulated by apoptotic signaling^19^. PAC-1 is a small-molecule drug that was initially developed to directly catalyze the conversion of caspase-3 from its inactive to its active state, thereby driving apoptosis in cancer cells^20^. This compound has completed a Phase I clinical trial, in which it produced multiple RECIST-confirmed responses^21^. However, the trial did not identify a biomarker for selecting a sensitive patient population. We previously applied CRISPR-based gene targeting to examine the reported MOA of PAC-1 and several other anti-cancer drugs^4^. We discovered that we could delete the gene encoding caspase-3 without impacting cellular sensitivity to PAC-1 treatment, indicating that PAC-1’s cytotoxic activity occurs through a caspase-3-independent mechanism.

The human genome encodes two executioner caspases that are paralogs of caspase-3: caspase-6 (CASP6) and caspase-7 (CASP7)^22^. These three proteases share overlapping substrates and are capable of functionally compensating for one another during apoptosis^18,23,24^. Previous research has suggested that PAC-1 could trigger cell death by activating caspase-6 or caspase-7 in addition to caspase-3^25,26^. We therefore sought to determine whether the redundant activation of multiple executioner caspases could explain PAC-1’s anti-cancer activity. We found that PAC-1 remains active in cells that lack all three executioner caspases. Instead, PAC-1 induces cancer cell death by depleting intracellular iron mediated by MDR1-driven efflux.

## Results

### The anti-cancer activity of PAC-1 does not require executioner caspase expression

In order to test the hypothesis that PAC-1’s anti-cancer activity results from the activation of alternative executioner caspases besides caspase-3, we generated a panel of cancer cell lines that lack executioner caspase expression. We used CRISPR/Cas9 to create triple caspase-3, caspase-6, and caspase-7 knockout clones (CASP3/6/7-KO) in A375 melanoma cells and HCT116 colorectal cancer cells (Figure 1A)^27^. As negative controls, we generated clones harboring gRNAs targeting the noncoding locus Rosa26. To verify complete target ablation, we confirmed the concurrent loss of executioner caspase expression using two independent antibodies for each protein (Figure 1B and S1A). We also obtained independently-derived CASP3/6/7-KO clones in the leukemia cell lines Jurkat and Nalm6, and we confirmed the absence of executioner caspase expression (Figure 1B)^24^. Active executioner caspases cleave PARP1 in response to DNA damage^28^. Accordingly, we treated cells with the topoisomerase poison doxorubicin and assessed PARP1 cleavage. Across all four cell lines, cleaved PARP was readily detectable in control cells but not in CASP3/6/7-KO clones, thereby further verifying total ablation of the executioner caspases (Figure S1B). Similarly, PAC-1 treatment resulted in PARP cleavage in wild-type cells but not in cells lacking caspase-3, -6, and -7 (Figure S1C).

**Figure 1.**
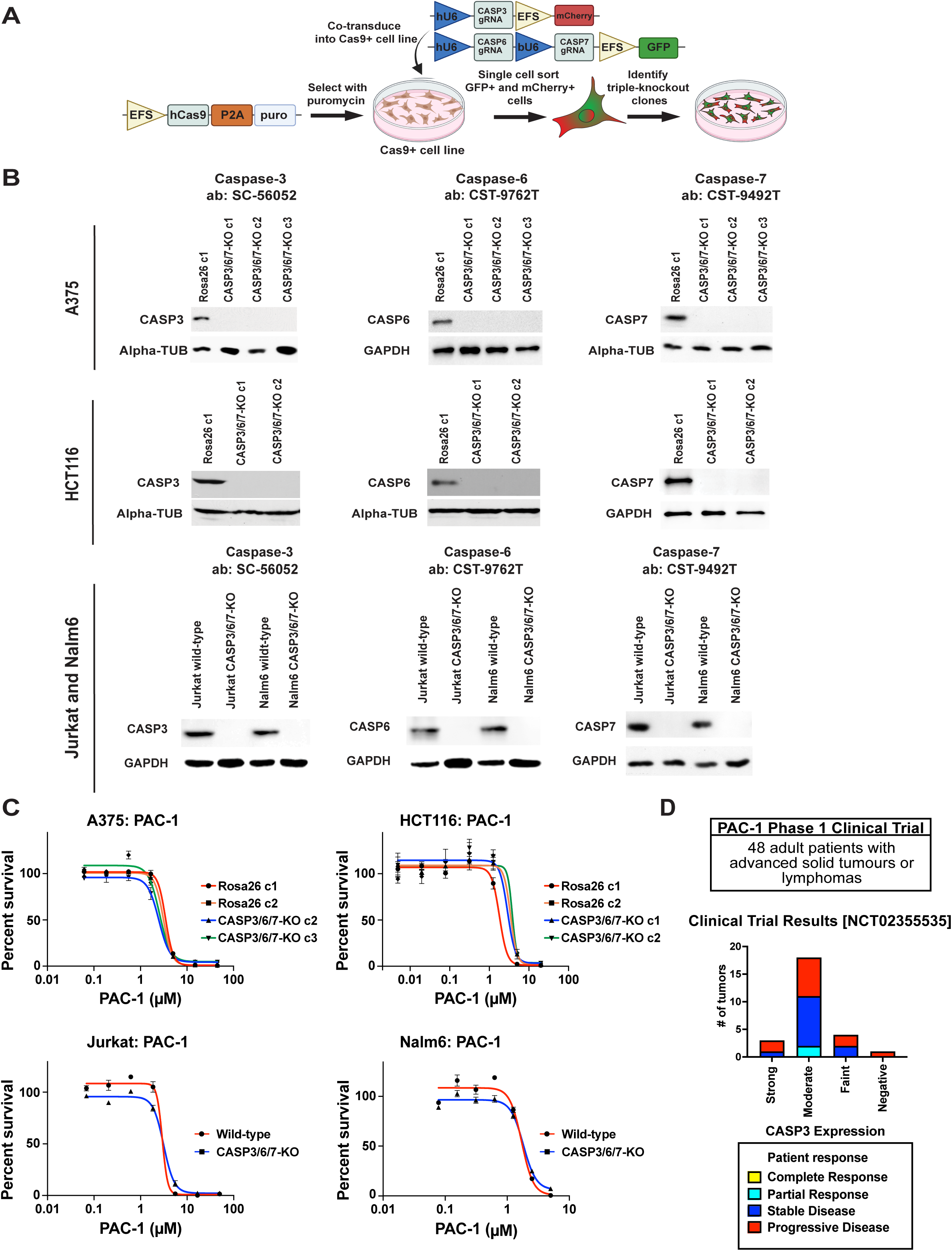
PAC-1 kills CASP3/6/7-knockout cancer cells. (A) Schematic of our approach to generate CASP3/6/7-knockout clones. (B) Western blot validation of the loss of caspase-3, caspase-6, and caspase-7 expression in the indicated cell lines. Alpha-tubulin or GAPDH levels were examined as a loading control. Additional western blots to verify caspase ablation are displayed in Figure S1. (C) 7-point dose response curves displaying cell viability in control or CASP3/6/7-KO clones exposed to the indicated concentrations of PAC-1. (D) A bar graph displaying patient response vs. caspase-3 expression in clinical trial NCT02355535.

Next, we performed drug sensitivity assays to test the viability of CASP3/6/7-expressing and CASP3/6/7-KO cells exposed to PAC-1. We found that there was no significant difference in PAC-1 toxicity between control and CASP3/6/7-KO cells in any of the four tested cancer cell lines (Figure 1C). For instance, PAC-1 exhibited a mean IC50 value of 3.3 μM in A375 Rosa26 clones, compared to a mean IC50 value of 2.9 μM in A375 CASP3/6/7-KO clones. These results indicate that PAC-1’s anti-cancer activity occurs independently of the executioner caspase pathway.

Recently, the results of the first clinical trial using PAC-1 have been published^21^. In this trial, two patients exhibited a RECIST-confirmed partial response and several exhibited long-term stable disease. However, caspase expression in the patients’ tumors was not reported. We obtained caspase-3 immunohistochemistry data for the tumors from this trial. Both responders exhibited moderate rather than strong caspase-3 expression, and across the entire cohort we did not observe any correlation between patient response and caspase-3 levels (Figure 1D). These results are consistent with our CRISPR-based cell culture experiments and indicate that caspase expression is unlikely to be an informative biomarker for PAC-1 sensitivity.

### PAC-1 treatment starves cancer cells of iron

In order to uncover PAC-1’s true MOA, we assessed drug screening data from the Broad Institute’s PRISM project^29^. This dataset is comprised of cell viability measurements for 537 cancer cell lines treated with 4,677 chemical compounds, including PAC-1. We calculated pairwise correlations between the vector of cellular sensitivities to PAC-1 and every other drug included in the PRISM screen. Surprisingly, PAC-1 exhibited the strongest correlation with deferasirox, a well-characterized iron chelation agent (Figure 2A)^30^. Cancer cell lines that were sensitive to deferasirox treatment also tended to be sensitive to PAC-1 treatment, and cancer cell lines that were resistant to deferasirox tended to be resistant to PAC-1 (R = 0.449, P < .0001; Figure 2B). These results raised the possibility that PAC-1 could function by interfering with cellular iron homeostasis rather than by activating caspases.

**Figure 2.**
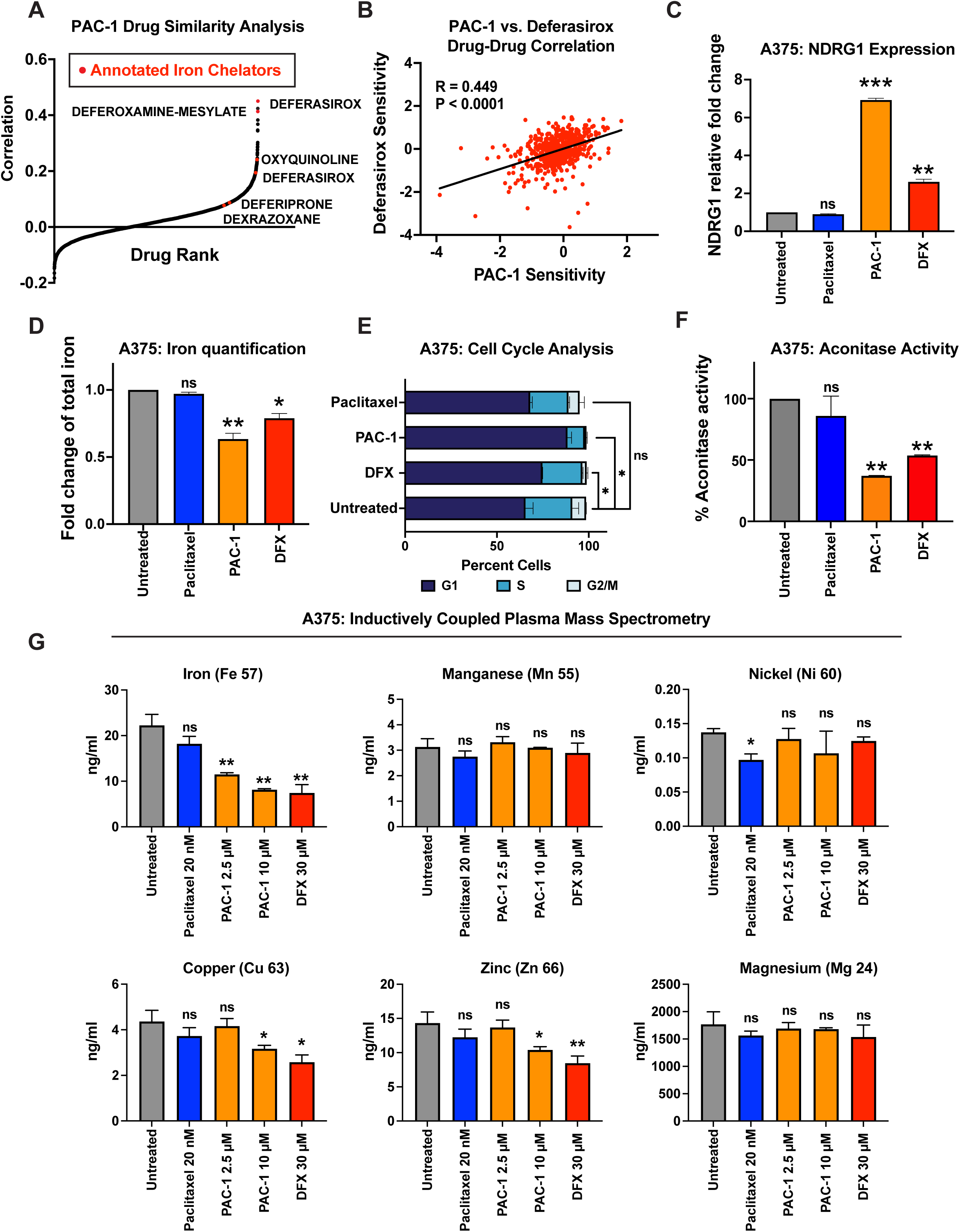
PAC-1 treatment causes iron starvation in cancer cells. (A) A waterfall plot displaying the correlation between the sensitivity profile of PAC-1 and every other drug included in the PRISM dataset^29^. Drugs annotated as iron chelators are highlighted with red dots. Note that deferasirox was run twice in this assay. (B) A scatterplot showing the correlation between sensitivity to deferasirox and sensitivity to PAC-1 in the PRISM dataset. The black line represents a linear regression plotted against the data. (C) Expression of the iron-starvation gene NDRG1 in A375 cells treated with 30 μM deferasirox (DFX), 10 μM PAC-1, or 20 nM paclitaxel. (D) Quantification of cellular iron in A375 cells treated with 30 μM deferasirox (DFX), 10 μM PAC-1, or 20 nM paclitaxel. (E) Cell cycle analysis in A375 cells treated with 30 μM deferasirox (DFX), 10 μM PAC-1, or 20 nM paclitaxel. (F) Quantification of aconitase activity in A375 cells treated with 30 μM deferasirox (DFX), 10 μM PAC-1, or 20 nM paclitaxel. (G) Quantification of the indicated metals in A375 cells treated with the indicated compounds and analyzed via ICP-MS. *, p < .05; **, p < .005; ***, p < .0005; Student’s t-test (two-sided).

To test whether PAC-1’s anti-cancer activity results from iron chelation, we conducted several complementary experiments. First, we treated A375 cells with PAC-1 and performed qPCR for the iron starvation gene NDRG1^31^. We found that exposure to PAC-1 or deferasirox resulted in a significant up-regulation of NDRG1 expression (Figure 2C). In contrast, treatment with paclitaxel, a widely used chemotherapy agent that targets microtubules, had no effect on the expression of NDRG1. Next, we directly quantified iron levels in cancer cells, and we observed that PAC-1 and deferasirox, but not paclitaxel, resulted in a significant decrease in cellular iron (Figure 2D).

Iron is an essential cofactor for the TCA cycle enzyme aconitase and for ribonucleotide reductase, which is required for nucleotide synthesis and entry into S phase^32,33^. We found that treatment with PAC-1 and deferasirox, but not paclitaxel, resulted in a G1 cell cycle arrest and a significant decrease in aconitase activity (Figure 2E-F). Finally, we tested the ability of excess iron to suppress PAC-1’s anti-cancer activity. We observed that 150 µM FeCl_2_ restored viability in both CASP3/6/7-expressing and CASP3/6/7-KO cells treated with either deferasirox or PAC-1 (Figure S2). In contrast, FeCl_2_ did not rescue the viability of cancer cells treated with paclitaxel. In total, these findings are consistent with our PRISM screening analysis and indicate that PAC-1 functions by starving cancer cells of iron.

### Cytotoxic concentrations of PAC-1 deplete cellular iron without affecting other metals

Many chelators exhibit limited specificity for individual metals and instead bind promiscuously to different metal ions^34^. Indeed, it has previously been suggested that PAC-1’s anti-cancer activity could result from its ability to relieve zinc-mediated repression of caspase activity^25^. We therefore sought to investigate whether PAC-1’s cytotoxicity could be specifically attributed to iron depletion or whether it instead resulted from interactions with additional metals. We treated A375 and HCT116 cells with 2.5 µM PAC-1, 10 µM PAC-1, 20 nM paclitaxel, or 5 µM of the zinc chelator TPEN and then quantified iron, zinc, and calcium levels via colorimetric assays. We observed that 2.5 µM PAC-1 was broadly cytotoxic and resulted in a 32% and 51% decrease in cellular viability in A375 and HCT116 cells, respectively (Figure S3A-B). We found that both concentrations of PAC-1 resulted in a significant decrease in cellular iron while neither concentration resulted in a significant decrease in cellular calcium (Figure S3C-F). Interestingly, 2.5 µM PAC-1 did not affect cellular zinc levels while 10 µM PAC-1 resulted in a moderate ∼10% decrease in zinc (Figure S3G-H). These experiments suggest that concentrations of PAC-1 that result in cell death affect iron levels without significantly depleting other metals, though supralethal concentrations of PAC-1 do deplete cellular zinc.

Next, we tested the ability of various concentrations of iron and zinc to restore viability in cells treated with PAC-1. We observed that 25 µM FeCl_2_ was sufficient to revert PAC-1-induced lethality in both A375 and HCT116 cells (Figure S3I-J). In contrast, while 25 µM ZnSO_4_ rescued the effects of TPEN on cell viability, it did not rescue the lethality of PAC-1. 100 µM ZnSO_4_ did partially mitigate PAC-1 cytotoxicity, though to a lesser extent than 100 µM FeCl_2_.

Finally, to assess the effects of PAC-1 on metal depletion in an unbiased manner, we performed inductively coupled plasma mass spectrometry (ICP-MS) to quantify metal abundance in A375 cells treated with PAC-1, TPEN, or paclitaxel. We found that 2.5 µM PAC-1 effectively depleted cellular iron but not cellular zinc, magnesium, manganese, copper, or nickel (Figure 2G). 10 µM PAC-1 resulted in a moderate decrease in cellular zinc and copper, though this was notably weaker than the effect of either concentration of PAC-1 on iron. In total, these complementary assays demonstrate that cytotoxic concentrations of PAC-1 deprive cells of iron without affecting the levels of several other biologically important metals, thereby implicating iron depletion as the key mechanism for PAC-1 lethality.

### Dual tridentate and bidentate binding of PAC-1 to iron

Spectroscopic chelation assays have previously shown that PAC-1 is capable of binding to Fe^2+^, but the stoichiometry and binding mode have not been identified^35^. We therefore determined the structure of a 1:1 complex of PAC-1 and iron by X-ray crystallography (Figure 3A, S4A and Table S1). The crystallographic structure reveals a remarkable decamer of PAC-1 and Fe^2+^ in a 1:1 ratio; though the structure in solution may differ, the stoichiometry agrees with the spectroscopic assessment below. Reacting two equivalents of PAC-1 with FeCl_2_ in a methanol/ethanol mixture gave a red crystalline solid in 85% yield (relative to Fe). The crystallographic data indicate that the piperazine group of PAC-1 is protonated, thus it has acted as a Brønsted base toward another molecule of PAC-1, which is deprotonated at the phenol group and hydrazone. Chloride ions are also present, supporting that basic functional groups are protonated. In the crystal structure, electron density adjacent to the N–benzyl nitrogen of each piperazine suggests that the piperazine moieties are protonated; the final model includes hydrogens in these locations.

**Figure 3.**
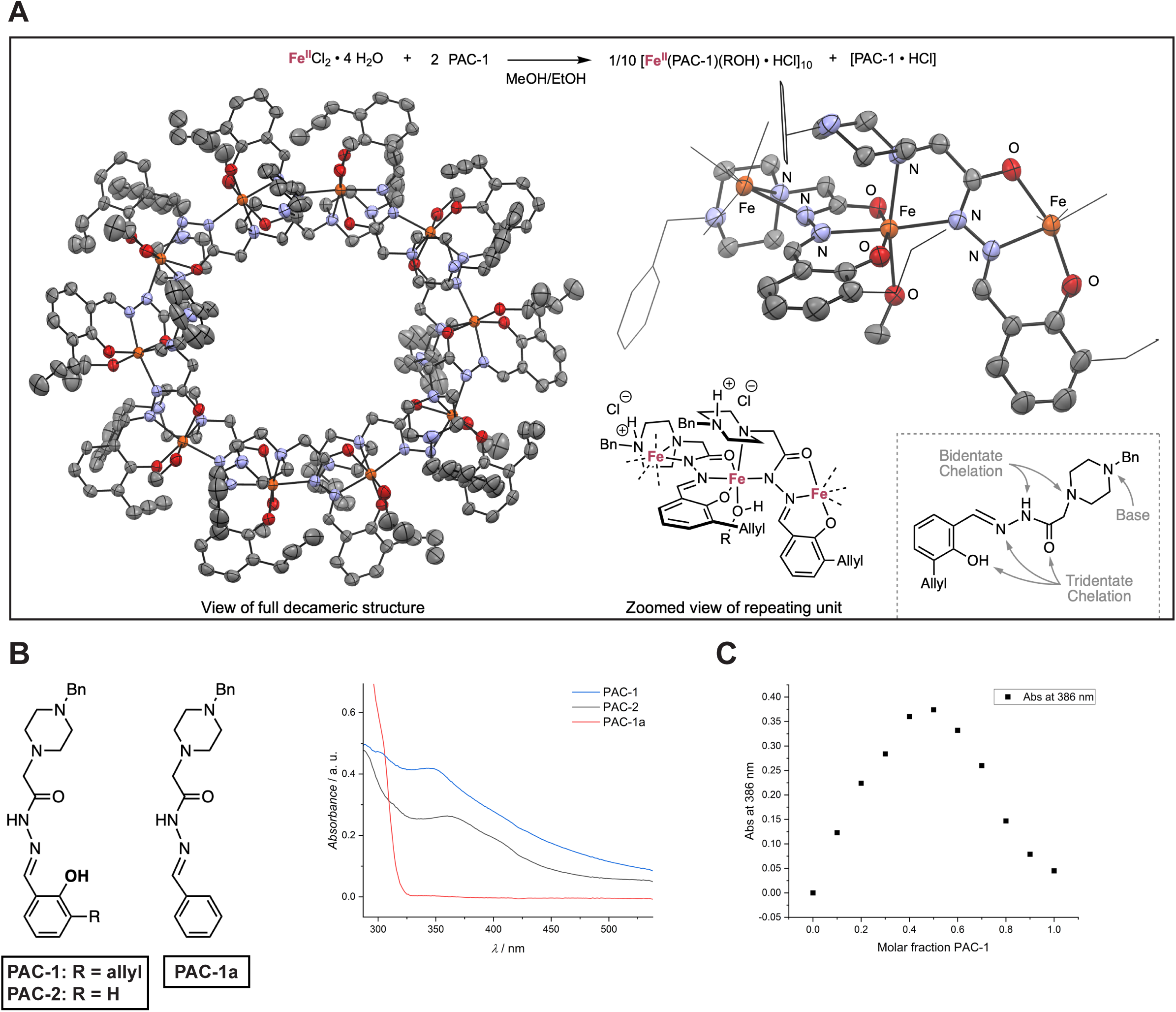
Structure of PAC-1 bound to iron. (A) Structure of the Fe^2+^ complex of PAC-1, crystallized from a solution of MeOH, EtOH, and toluene. The left shows a plot of the full decameric structure and the right shows a zoomed view of the repeating and structural diagram (R = Me or Et). The ORTEP diagrams show 50% ellipsoids with hydrogens and chlorides omitted for clarity. The dashed box contains a diagram of the metal/proton binding sites. (B) UV/Vis spectra of PAC-1, PAC-1a and PAC-2 (50 µM each) in the presence of 50 µM FeCl_2_ in 50 mM HEPES buffer, pH = 7.4. (C) The absorbance for solutions of PAC-1:Fe^2+^ mixtures in buffered solution (50 mM HEPES, pH = 7.4). The maximum at 1:1 indicates the stoichiometry of the complex.

Based on the crystal structure, we hypothesized that PAC-1’s hydroxyl group would be essential for iron binding and the allyl group would be dispensable. To test this prediction, we studied the chelation of Fe^2+^ with PAC-1 and two related analogues: PAC-1a, in which the hydroxyl and allyl groups are replaced with hydrogens, and PAC-2, in which only the allyl group is replaced with a hydrogen (Figure 3B). In 50 mM HEPES buffer at pH 7.4, both PAC-1 and PAC-2 chelate Fe^2+^, as indicated by the broad bands between 300 and 500 nm (Figure 3B and S4B). In contrast, PAC-1a shows no spectroscopic changes that indicate binding (Figure 3B). This suggests that the hydroxyl group enables tridentate iron chelation. Next, we used the method of continuous variation to verify that the binding stoichiometry was 1:1 between PAC-1 and Fe^2+^ (Figure 3C). 1:1 binding is also consistent with ^1^H NMR and Mössbauer spectra analysis (Figure S4C-E).

While the decameric structure of PAC-1 bound to iron that we characterized in the solid state may break up in solution, it shows clearly that PAC-1 can coordinate to Fe^2+^ in a tridentate mode (O,N,O via the phenol and hydrazide) and a bidentate mode (N,N via the hydrazide and piperazine). The piperazine moiety of PAC-1 enables this coordination. In total, the complex demonstrates that (1) PAC-1 can readily coordinate to Fe^2+^, (2) the piperazine moiety enables an additional chelation site for metal ions, and (3) the nitrogen(s) of the piperazine can act as Brønsted bases and may be protonated under certain biological conditions. Our results therefore establish a structural basis for the interaction of PAC-1 and Fe^2+^ ions.

### PAC-1 is selectively toxic towards MDR1-overexpressing cancer cells

As our data demonstrated that the cellular response to PAC-1 does not depend on the executioner caspases, we sought to discover an alternative biomarker capable of predicting PAC-1 sensitivity. We compared the effects of PAC-1 exposure on cellular viability with mutation, copy number, and gene expression data from every cancer cell line included in the PRISM dataset. Surprisingly, the strongest predictor of sensitivity to PAC-1 was high expression of the gene ABCB1 (Figure 4A and S5A). ABCB1 encodes the efflux pump MDR1, and high MDR1 expression causes drug resistance and chemotherapy failure^12^. However, our findings unexpectedly indicated that MDR1 expression promotes sensitivity, rather than resistance, to PAC-1 treatment.

**Figure 4.**
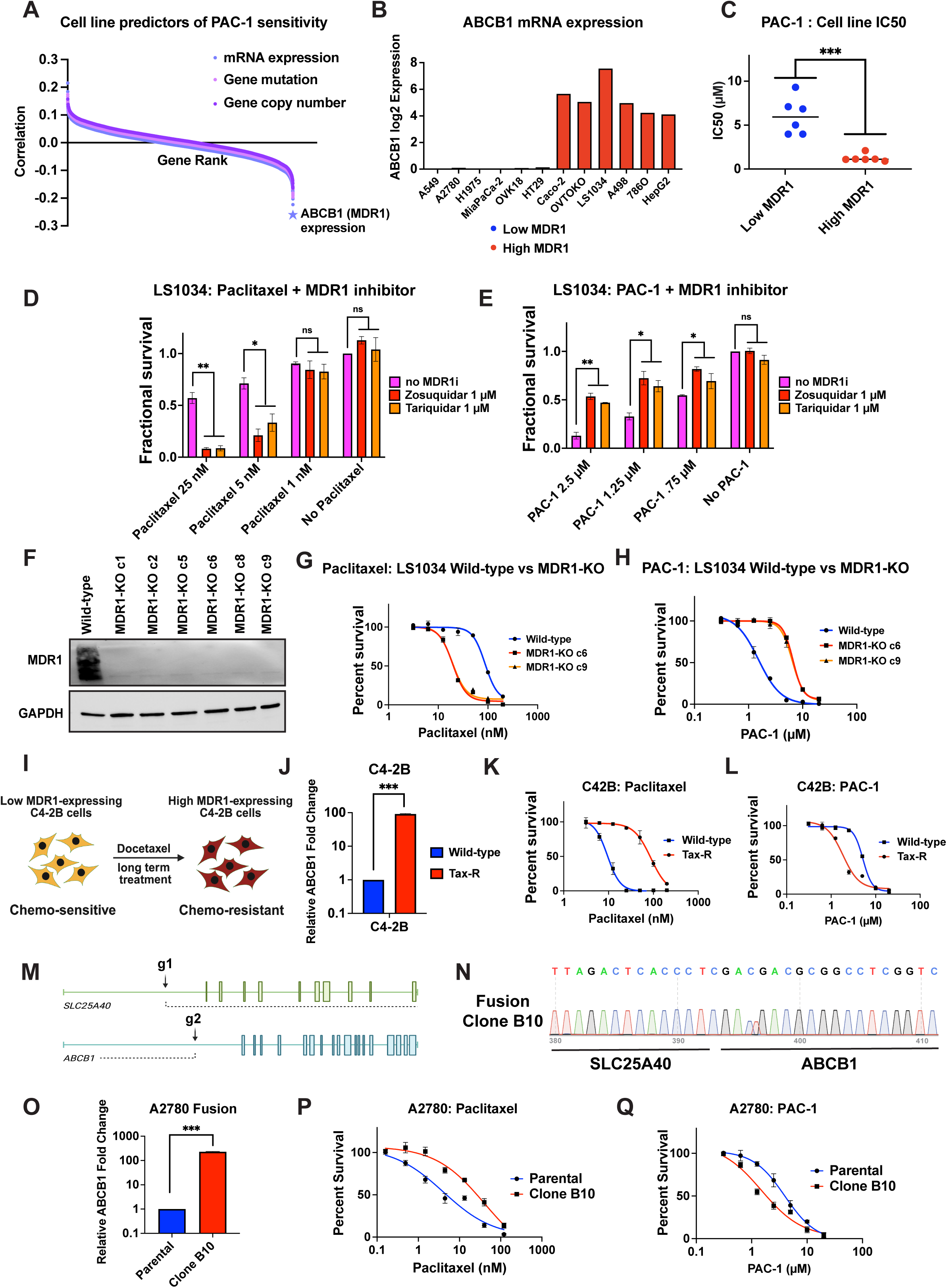
MDR1 expression increases sensitivity to PAC-1. (A) A waterfall plot displaying the correlation between gene expression, mutation, and copy number and PAC-1 sensitivity in the PRISM dataset^29^. The expression of ABCB1 (MDR1) is highlighted with a star. (B) ABCB1 expression levels in 12 different cancer cell lines are displayed^84^. (C) The IC50 values of PAC-1 across the 12 cell lines included in (B) are displayed. (D) A bar graph displaying the viability of MDR1-expressing LS1034 cells treated with varying concentrations of paclitaxel in the presence or absence of two MDR1 inhibitors, zosuquidar and tariquidar. (E) A bar graph displaying the viability of MDR1-expressing LS1034 cells treated with varying concentrations of PAC-1 in the presence or absence of two MDR1 inhibitors, zosuquidar and tariquidar. (F) Western blot validation of the loss of MDR1 expression in LS1034 cells. GAPDH levels were examined as a loading control. (G) 7-point dose response curves displaying cell viability in LS1034 wild-type or MDR1-KO clones exposed to the indicated concentrations of paclitaxel. (H) 7-point dose response curves displaying cell viability in LS1034 wild-type or MDR1-KO clones exposed to the indicated concentrations of PAC-1. (I) A schematic displaying the isolation of docetaxel-resistant, MDR1-expressing C4-2B prostate cancer cells^39^. (J) Expression of ABCB1 in wild-type and docetaxel-resistant C4-2B (Tax-R) cells. (K) 7-point dose response curves displaying cell viability in wild-type or docetaxel-resistant C4-2B (Tax-R) cells exposed to the indicated concentrations of paclitaxel. (L) 7-point dose response curves displaying cell viability in wild-type or docetaxel-resistant C4-2B (Tax-R) cells exposed to the indicated concentrations of PAC-1. (M) Structure of the SLC25A40 and ABCB1 loci on chromosome 7 and the sites targeted by gRNAs designed to produce an SLC25A40-ABCB1 fusion. (N) Sequence verification of an SLC25A40-ABCB1 fusion clone (B10) in A2780 cells. (O) Expression of ABCB1 in wild-type A2780 cells and in A2780 cells harboring an SLC25A40-ABCB1 fusion. (P) 7-point dose response curves displaying cell viability in wild-type A2780 cells and in A2780 cells harboring an SLC25A40-ABCB1 fusion exposed to the indicated concentrations of paclitaxel. (Q) 7-point dose response curves displaying cell viability in wild-type A2780 cells and in A2780 cells harboring an SLC25A40-ABCB1 fusion exposed to the indicated concentrations of PAC-1. *, p < .05; **, p < .005; ***, p < .0005; Student’s t-test (two-sided).

To validate the results of this analysis, we assessed the effects of PAC-1 across a panel of cancer cell lines that express high levels or low levels of MDR1 (Figure 4B). We found that the median IC50 value for PAC-1 was 1.2 μM across MDR1-high cell lines and 6.0 μM across MDR1-low cell lines (Figure 4C). Next, we sought to determine whether MDR1 activity directly increases sensitivity to PAC-1. We exposed LS1034 cells, a colorectal cancer cell line that expresses high levels of MDR1, to two well-characterized MDR1 inhibitors, zosuquidar and tariquidar^36^. Paclitaxel is an established MDR1 substrate, and, as expected, treatment with the MDR1 inhibitors increased the sensitivity of LS1034 cells to paclitaxel (Figure 4D)^37^. In contrast, treatment with zosuquidar and tariquidar resulted in a significant decrease in sensitivity to PAC-1, implicating MDR1 activity as a cause of PAC-1-induced lethality (Figure 4E). Finally, we used CRISPR to delete the ABCB1 gene from LS1034 cells, and we examined the effects of drug treatment on isogenic MDR1-expressing and MDR1-knockout cells (Figure 4F). We found that the MDR1-KO clones were 5-fold more sensitive to paclitaxel compared with the MDR1-expressing parental cell line, but these same knockout clones were 4-fold less sensitive to PAC-1 (Figure 4G-H).

Next, we sought to explore the relationship between MDR1 expression and PAC-1 sensitivity in two clinically-relevant contexts. First, cancer treatment with a variety of chemotherapy agents results in MDR1 upregulation and drug resistance^38^. To model this process, we assessed C4-2B prostate cancer cells that had been exposed to docetaxel for nine months (Figure 4I)^39^. qPCR analysis revealed that the evolved cell population exhibited a 91-fold increase in ABCB1 expression relative to the parental cell line (Figure 4J). While these evolved cells displayed a significant decrease in sensitivity to paclitaxel, they were 3-fold more sensitive to PAC-1 compared with the paclitaxel-responsive parental cell population (Figure 4K-L). Second, recent genomic sequencing efforts have demonstrated that MDR1 activation in the context of recurrent disease commonly occurs through the acquisition of intrachromosomal fusions that place ABCB1 under the control of highly-expressed promoters near the ABCB1 locus. The most frequently-observed fusion in a cohort of patients with drug-resistant ovarian and breast cancer was between ABCB1 and the SLC25A40 gene^14,40^. We used CRISPR to generate this fusion event in the MDR1-low ovarian cancer cell line A2780 (Figure 4M-N). Fusion-positive cells exhibited a 230-fold increase in ABCB1 expression, a 9-fold decrease in paclitaxel sensitivity, and a 3-fold increase in PAC-1 sensitivity (Figure 4O-Q). In total, our data demonstrate that, across a variety of cell lines and model systems, high MDR1 expression enhances cellular sensitivity to PAC-1 treatment.

### PAC-1 sensitivity is not influenced by the expression of alternative drug efflux transporters

In addition to MDR1, several other cellular transporters have been demonstrated to contribute to chemotherapy resistance. The best-characterized of these alternative drug efflux pumps include ABCC1, which encodes MRP1, and ABCG2, which encodes BCRP^41^. We sought to determine whether the expression of these transporter genes could also influence PAC-1 sensitivity. First, we analyzed the expression of these genes in the PRISM dataset, and we found that there was no significant correlation between the expression of either ABCC1 or ABCG2 and sensitivity to PAC-1 across cancer cell lines (Figure S5B-C). Next, we ectopically overexpressed ABCB1, ABCC1, or ABCG2 in MiaPaCa-2 pancreas cancer cells, which normally express low levels of all three genes (Figure S5D). Overexpressing ABCB1 significantly increased the sensitivity of MiaPaCa-2 cells to PAC-1, while overexpressing either ABCC1 or ABCG2 had no effect on PAC-1 sensitivity (Figure S5E-H). In total, our data indicate that PAC-1 is selectively toxic towards cancer cells that overexpress MDR1 but not other drug efflux pumps.

### PAC-1 is active against MDR1-overexpressing patient-derived ovarian cancer cells

We sought to determine whether PAC-1’s activity against MDR1-expressing cells extends beyond established cancer cell lines. Toward that goal, we investigated a panel of *ex vivo* ovarian cancer models (OCMs) that were derived from patient ascites with minimal culturing (Figure S6A)^42,43^. Gene expression analysis revealed a wide range of ABCB1 expression across the OCMs (Figure S6B). Unlike the established cancer cell lines, some of which did not express any ABCB1, all OCMs expressed detectable levels of ABCB1, though there was a 500-fold difference in ABCB1 expression between the lowest OCM (OCM.118-7) and the highest OCM (OCM-149). Next, we tested the effects of PAC-1 exposure on colony formation in each OCM. Consistent with the cell line results described above, we found that OCMs expressing high levels of ABCB1 were significantly more sensitive to PAC-1 than OCMs expressing low or moderate levels of ABCB1 (P < .03; Figure S6C-D). Indeed, the four OCMs that were most sensitive to PAC-1 also expressed the highest levels of ABCB1, and there was a significant correlation between ABCB1 expression and PAC-1 sensitivity across the entire OCM panel (Rho = -0.883; P < .004; Figure S6D). We conclude that PAC-1 is capable of eliminating both established cancer cell lines and minimally passaged patient-derived ovarian cancer cells that express high levels of MDR1.

### The activity of PAC-1 against MDR1-high cells is unique compared with approved cancer drugs

We next sought to discover whether other compounds with mechanisms similar to PAC-1 were capable of eliminating cells that overexpress MDR1. The small-molecule drug 1541B has been reported to function like PAC-1 by directly activating executioner caspases^44^. As with PAC-1, we found that its anti-cancer activity was not affected by the deletion of all executioner caspases, indicating that it functions via some other mechanism (Figure S7A). However, 1541B exhibited equivalent potency against MDR1-WT and MDR1-KO LS1034 cells (Figure S7B). This demonstrates that MDR1-selectivity is not a shared property of compounds developed to activate the executioner caspases. Additionally, we found that the MDR1 inhibitor zosuquidar does not exhibit MDR1-selective cell killing, indicating that blocking MDR1-mediated efflux is unlikely to be the mechanism driving MDR1-specific toxicity (Figure S7C). Similarly, the iron chelator deferasirox and the zinc chelator TPEN were equally effective against MDR1-WT and MDR1-KO cells (Figure S7D-E). Thus, compounds that share potential targets and mechanisms with PAC-1 do not necessarily share its ability to selectively eliminate MDR1-expressing cancer cells.

Next, we tested whether other drugs that are marketed or in development for cancer therapy were able to target cells overexpressing MDR1. We screened 130 cancer drugs in the MDR1-WT and MDR1-KO LS1034 cells, and we identified 73 compounds that were cytotoxic in this cell line at concentrations below 20 µM (Figure S8 and Table S2). As expected, MDR1 expression decreased sensitivity to multiple agents, including selumetinib, topotecan, and dactinomycin. Within this panel, the only compound for which MDR1 expression increased sensitivity was PAC-1 (Figure S8 and Table S2). These results indicate that PAC-1’s selective activity against MDR1-expressing cells is a unique property not broadly shared by other anti-cancer agents in clinical testing.

### PAC-1 is an MDR1 substrate whose efflux is enhanced by iron

We sought to uncover the mechanism by which MDR1 expression promotes PAC-1 sensitivity. Toward this goal, we investigated whether PAC-1 is subject to MDR1-mediated efflux. The fluorescent dye rhodamine-123 is effluxed from cells in an MDR1-dependent process, and when cells transiently incubated with rhodamine-123 are exposed to a competing MDR1 substrate, rhodamine-123 efflux is delayed (Figure 5A)^45^. While wild-type LS1034 cells rapidly lost rhodamine-123 fluorescence when moved to rhodamine-free media, rhodamine-123 efflux was decreased by either exposure to paclitaxel or the deletion of ABCB1 (Figure 5B). PAC-1 treatment also resulted in a dose-dependent delay in rhodamine-123 efflux, though at higher concentrations than required for paclitaxel. Next, we quantified the ATPase activity of purified MDR1, which is stimulated upon binding to MDR1 substrates^46^. Exposure to PAC-1 significantly enhanced MDR1’s ATPase activity, albeit to a lesser extent than the well-characterized MDR1 substrate verapamil (Figure 5C-D). Taken together, these experiments indicate that PAC-1 is indeed an MDR1 substrate, though they suggest that PAC-1 may be a weaker substrate than canonical MDR1 substrates like paclitaxel and verapamil.

**Figure 5.**
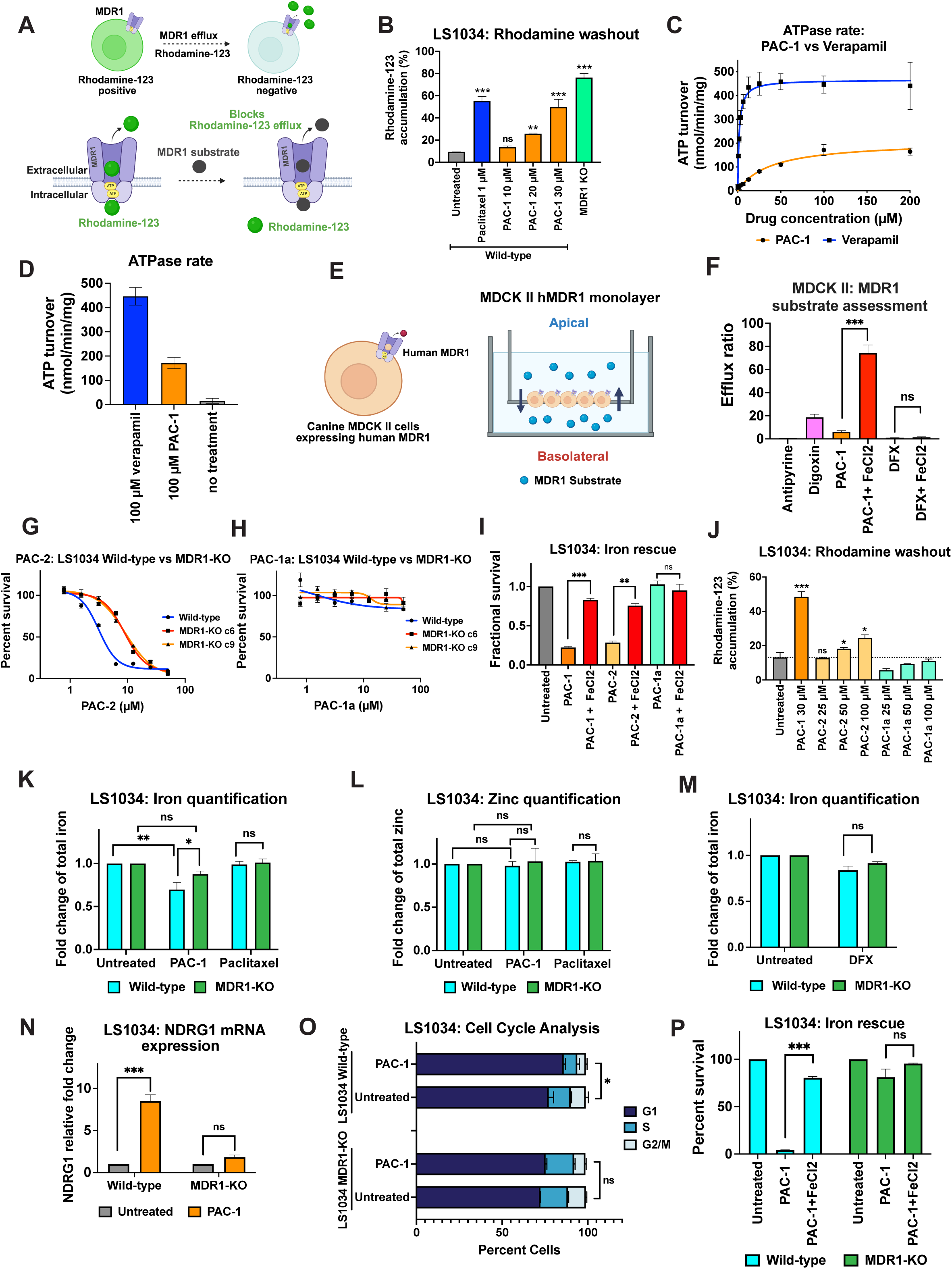
MDR1 enhances PAC-1-mediated iron depletion. (A) A schematic of the rhodamine-123 efflux assay for quantifying MDR1-drug interactions. (B) A bar graph displaying the percent of rhodamine-positive LS1034 cells, one hour after rhodamine washout, in the presence of the indicated compounds or in LS1034 MDR1-KO cells. (C) MDR1 ATPase activity as a function of PAC-1 and verapamil concentration. Data represent means and SEMs from three technical replicates of three biological replicates (n=9) collected at 37°C. (D) MDR1 ATPase activity in the presence of 100 μM PAC-1 or 100 μM verapamil concentration. (E) A schematic of the MDCK-II monolayer assay for quantifying MDR1-mediated efflux. (F) A bar graph displaying the MDR1 efflux ratio of 10 μM of the indicated compounds, in the presence or absence of excess iron. Antipyrine is not effluxed by MDR1 and serves as a negative control while digoxin is a recognized MDR1 substrate and serves as a positive control. (G) 7-point dose response curves displaying cell viability in LS1034 wild-type or MDR1-KO clones exposed to the indicated concentrations of PAC-2. (H) 7-point dose response curves displaying cell viability in LS1034 wild-type or MDR1-KO clones exposed to the indicated concentrations of PAC-1a. (I) Cellular viability in LS1034 cells treated with 2.5 μM PAC-1, 5 μM PAC-2, or 20 μM PAC-1a, in the presence or absence of 150 µM FeCl_2_. (J) A bar graph displaying the percent of rhodamine-positive LS1034 wild-type cells, one hour after rhodamine washout, in the presence of the indicated compounds. (K) Quantification of cellular iron in LS1034 wild-type and MDR1-KO cells treated with 10 μM PAC-1 or 20 nM paclitaxel. (L) Quantification of cellular zinc in LS1034 wild-type and MDR1-KO cells treated with 10 μM PAC-1 or 20 nM paclitaxel. (M) Quantification of cellular iron in LS1034 wild-type and MDR1-KO cells treated with 10 μM deferasirox. (N) Expression of the iron-starvation gene NDRG1 in LS1034 wild-type and MDR1-KO cells treated with 10 μM PAC-1. (O) Cell cycle analysis in LS1034 wild-type and MDR1-KO cells treated with 10 μM PAC-1. (P) Cellular viability in LS1034 wild-type and MDR1-KO cells treated with 2.5 μM PAC-1, in the presence or absence of 150 µM FeCl_2_. *, p < .05; **, p < .005; ***, p < .0005; Student’s t-test (two-sided).

To further verify these observations, we quantitatively assessed MDR1-mediated drug efflux across a monolayer of canine MDCK-II cells expressing human MDR1 (Figure 5E)^47^. Under standard culture conditions, PAC-1 exhibited an efflux ratio of 6, while the established MDR1 substrate digoxin^48^ exhibited an efflux ratio of 19, indicating that PAC-1 is subject to a moderate degree of MDR1-driven efflux (Figure 5F). Remarkably, in the presence of excess iron, PAC-1’s MDR1 efflux increased to 74 (P < .001). In contrast, the addition of iron did not affect the efflux of deferasirox. Together, these data demonstrate that PAC-1 is a moderate MDR1 substrate, and the presence of iron dramatically enhances the MDR1-mediated efflux of PAC-1 but not other iron chelators.

### A PAC-1 analog that lacks iron-binding activity does not exhibit MDR1-selective cell killing

Based on our cellular and biochemical assessment of PAC-1 activity, we hypothesized that the MDR1-selective cytotoxicity displayed by PAC-1 is driven by the MDR1-mediated efflux of PAC-1 bound to cellular iron. If this theory is accurate, then the ability of PAC-1 to bind iron should be essential for its MDR1-selective cytotoxicity. To test this hypothesis, we examined the two PAC-1 analogs that we had previously profiled: PAC-1a, which lacks the ability to bind iron, and PAC-2, which maintains iron-binding activity (Figure 3B). We found that PAC-2 exhibited greater toxicity in MDR1-expressing cells compared with MDR1-KO cells, though its selectivity and potency were moderately lower than PAC-1 (Figure 5G). In contrast, PAC-1a exhibited no cytotoxicity in either MDR1-expressing or MDR1-KO cells at concentrations up to 50 μM (Figure 5H). PAC-2-mediated lethality could be reversed by iron supplementation, and PAC-2, but not PAC-1a, moderately delayed rhodamine-123 efflux (Figure 5I-J). These results are consistent with our hypothesis that MDR1-selective cell killing by PAC-1 is driven by PAC-1’s ability to deplete cellular iron in an MDR1-dependent manner.

### MDR1 enhances PAC-1-mediated iron starvation

Next, we characterized the impact of MDR1 expression on iron homeostasis in cancer cells treated with PAC-1. We found that exposure to PAC-1 resulted in a significantly greater decrease in cellular iron in MDR1-expressing cells compared with MDR1-KO cells (Figure 5K). In contrast, this same concentration of PAC-1 had no effect on cellular zinc in either MDR1-expressing or MDR1-KO LS1034 cells (Figure 5L). Deferasirox treatment resulted in a similar decrease in iron levels in both wild-type and MDR1-KO cells, thereby providing a rationale as to why this iron chelator does not exhibit MDR1-selective cell killing (Figure 5M). PAC-1 treatment caused the upregulation of the iron starvation gene NDRG1 and a G1 cell cycle arrest in wild-type cells, but these effects were lost in the absence of MDR1 (Figure 5N-O). Finally, iron supplementation rescued PAC-1 lethality in MDR1-expressing but not MDR1-KO cells (Figure 5P). In total, our data indicate that PAC-1 is an MDR1 substrate, and MDR1 enhances PAC-1-mediated iron depletion in cancer cells. Our results are consistent with a model in which iron binding enhances the efflux of PAC-1 by MDR1, thereby starving cancer cells of iron over time.

### Leveraging PAC-1 to redirect cancer evolution towards a drug-sensitive state

MDR1 expression causes resistance to dozens of widely-used chemotherapy agents^49–51^. We hypothesized that we could take advantage of PAC-1’s selective activity against MDR1-overexpressing cells to drive cancers into an MDR1-low, drug-sensitive cellular state. Toward that goal, we labeled LS1034 MDR1-WT and MDR1-KO with different fluorophores, mixed the two populations together at equal concentrations, and then measured the relative abundance of each population over time (Figure 6A)^52^. In normal media, the two populations maintained a 50:50 ratio, indicating that MDR1 expression has minimal impact on cell fitness in the absence of drug treatment (Figure 6B). In the presence of paclitaxel, the MDR1-KO cells were rapidly outcompeted by the MDR1-WT cells. However, this pattern was reversed in the presence of PAC-1, and after three passages the MDR1-KO outnumbered the MDR1-WT cells by a 3:1 ratio. These experiments indicate that in a mixed population of chemotherapy-resistant and chemotherapy-sensitive cells, the chemotherapy-resistant cells can be eliminated by exposure to PAC-1 (Figure 6B).

**Figure 6.**
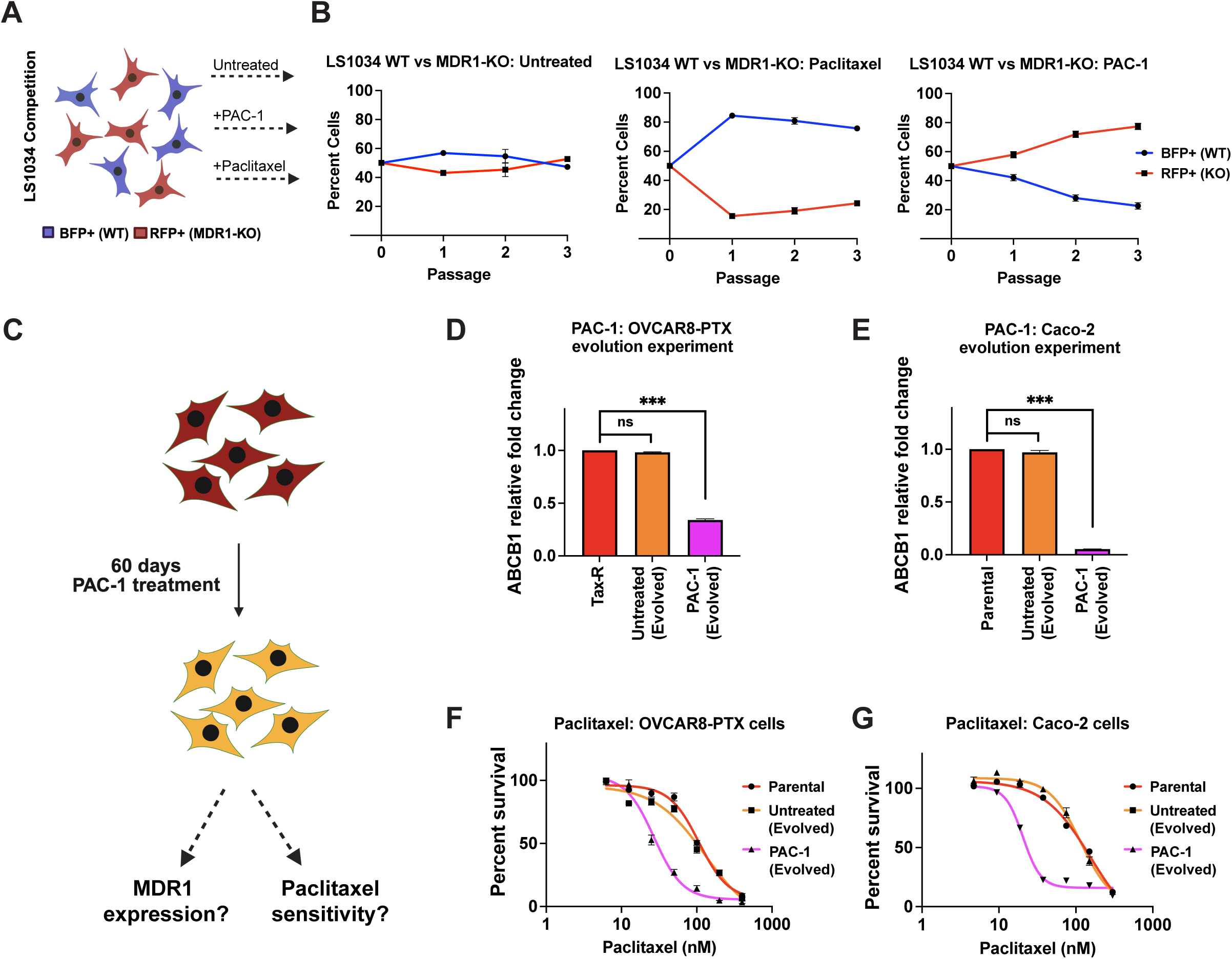
Restoring chemotherapy sensitivity via PAC-1 treatment. (A) Schematic of a cellular competition assay in which LS1034 wild-type and MDR1-KO cells are labeled with either BFP (WT) or RFP (MDR1-KO), mixed together at a 50-50 ratio, and then grown in the presence of different drugs. (B) Time courses displaying the relative abundance of BFP-expressing and RFP-expressing cells grown in normal media, in the presence of 10 nM paclitaxel, or in the presence of 3 μM PAC-1, measured every four days. (C) A schematic of the experimental approach to examine the effects of PAC-1 on ABCB1 expression and drug sensitivity in cancer cells. (D) qPCR analysis of ABCB1 expression in OVCAR8-PTX cells grown in normal media or grown in the presence of PAC-1 for 50 days. (E) qPCR analysis of ABCB1 expression in Caco-2 cells grown in normal media or grown in the presence of PAC-1 for 60 days. (F) 7-point dose response curves displaying cell viability in parental OVCAR8-PTX cells, OVCAR8-PTX cells grown in normal media, and OVCAR8-PTX cells grown in the presence of PAC-1, exposed to the indicated concentrations of paclitaxel. (G) 7-point dose response curves displaying cell viability in parental Caco-2 cells, Caco-2 cells grown in normal media, and Caco-2 cells grown in the presence of PAC-1, exposed to the indicated concentrations of paclitaxel.

Next, we sought to discover whether exposure to PAC-1 could be used to sensitize drug-resistant cells to chemotherapy treatment. Toward that goal, we tested the colorectal cancer cell line Caco-2 and a drug-resistant variant of the ovarian cancer cell line OVCAR8 (OVCAR8-PTX)^53^, both of which express high levels of MDR1. We cultured each cell line in either normal media or in a sub-lethal concentration of PAC-1 for 50-60 days, and then we assessed ABCB1 expression via qPCR (Figure 6C). While growth in normal media had no effect on ABCB1 expression, we found that exposure to PAC-1 treatment resulted in a significant downregulation of ABCB1 in both OVCAR8-PTX and Caco-2 cells (Figure 6D-E). Finally, we tested whether chronic PAC-1 exposure resulted in collateral sensitivity to paclitaxel. Indeed, we found that the evolved OVCAR8-PTX cells were 4-fold more sensitive to paclitaxel compared with the parental cell population and the evolved Caco-2 cells were 7-fold more sensitive to paclitaxel compared with the parental cell population (Figure 6F-G). We conclude that PAC-1 treatment consistently decreases MDR1 expression and restores sensitivity to chemotherapy agents in previously drug-resistant cells.

## Discussion

Here, we redefine the mechanism of action of the clinical-stage oncology compound PAC-1 and reveal a highly selective vulnerability in multidrug-resistant cancers. Contrary to its original purpose as a procaspase activator, PAC-1 induces cytotoxicity independently of the executioner caspases and instead functions as an iron chelator. We demonstrate that PAC-1 binds Fe²⁺ ions through a dual tridentate and bidentate coordination mode, effectively starving cancer cells of iron while sparing other biologically important metals. This iron depletion disrupts metabolic enzymes and DNA synthesis, triggering G1 arrest and cell death. Remarkably, PAC-1 exhibits potent and preferential activity against MDR1-overexpressing cancer cells, including chemotherapy-resistant and patient-derived models that are refractory to other agents. Our mechanistic studies show that MDR1 actively effluxes the iron-bound form of PAC-1, thereby accelerating cellular iron loss and driving the selective killing of drug-resistant cells. These results establish a powerful therapeutic paradigm in which MDR1 expression can be exploited as a collateral vulnerability in refractory malignancies through progressive iron depletion.

MDR1 expression is a pervasive cause of treatment failure in oncology. The MDR1 transporter expels a broad range of structurally-diverse compounds, including cytotoxic chemotherapies and targeted therapies as well as next-generation modalities like antibody-drug conjugates and PROTACs^54,55^. MDR1 expression prevents drug accumulation, effluxing anti-cancer agents before they can exert a cytotoxic effect. Previous efforts to combat MDR1-mediated drug resistance have focused on the development of competitive inhibitors to block MDR1 activity. While more than a dozen MDR1 inhibitors entered clinical trials, these drugs exhibited limited specificity for MDR1 relative to other important cellular transporters and resulted in unacceptable levels of toxicity^55,56^. Despite decades of research, no clinically approved strategy has succeeded in blocking MDR1-driven resistance, underscoring the need for alternative approaches that exploit, rather than inhibit, MDR1 function.

To date, directly targeting MDR1 as a potential cause of drug sensitivity has not been tested in cancer patients. Preclinical compound screens have been conducted to identify vulnerabilities specific for MDR1-overexpressing cancer cells^57,58^. These and similar efforts have identified an array of structurally-diverse compounds with various reported MOAs, including ROS overproduction^59^, STAT3 inhibition^60^, ATP depletion^61^, and survivin inhibition^62^, as well as many compounds with unknown or unclear cellular targets^29,63^. Interestingly, many drugs identified in these earlier efforts were observed to be metal chelators or complexes with different metals, including copper, tin, molybdenum, and lanthanum^57,64^. The mechanism(s) by which these compounds induced cell death was unclear, and it was speculated that they could carry toxic quantities of metal into MDR1-expressing cells^57^. One of these chelators was shown to target iron^65^, and we hypothesize that some of these drugs may exhibit an MOA similar to PAC-1 and induce selective iron depletion, rather than toxic metal accumulation, in an MDR1-dependent manner.

Among the various mechanisms that have been reported for targeting MDR1-expressing tumors, progressive iron depletion may be a particularly tractable therapeutic approach. Cancer cells require more iron than normal tissue, which may be necessary to sustain their elevated levels of ribonucleotide reductase activity, ATP production, and redox demands^66^. Iron chelation results in PARP cleavage, caspase activation, and apoptotic signaling, as observed in PAC-1-treated cells, which may be a consequence of DNA damage and replication fork collapse caused by ribonucleotide reductase inhibition^20,67–70^. Moreover, iron chelators like deferasirox have been used clinically for several decades to treat iron overload syndrome, and these agents have demonstrated favorable safety profiles and pharmacodynamic properties^30^. While deferasirox and other chelators have previously been tested as anti-cancer agents, their clinical activity has been modest, potentially because these trials were not guided by patient selection biomarkers and these compounds do not possess the powerful MOA reported here^66^. By coupling iron chelation to MDR1 efflux, PAC-1 converts a canonical mechanism of drug resistance into a driver of selective vulnerability, potentially establishing a path to therapeutically exploit multidrug resistance in the clinic.

Our results provide a potential explanation for the outcomes of the initial Phase I trial of PAC-1, which reported multiple durable responses but failed to identify a predictive biomarker based on the presumed mechanism of caspase activation^21^. The discovery that PAC-1 functions as an iron chelator whose activity is potentiated by MDR1 expression offers a compelling, alternative biomarker hypothesis. While our preclinical data across cell lines, drug-resistant models, and patient-derived OCMs consistently link high MDR1 expression to PAC-1 sensitivity, it remains to be seen whether MDR1 expression can effectively predict PAC-1 sensitivity in patients. Future clinical studies, guided by this new mechanistic framework, will be essential to validate MDR1 as a biomarker for selecting patients most likely to benefit from this therapeutic strategy, potentially transforming a common driver of drug resistance into a targetable clinical vulnerability.

## Acknowledgments

Research in the Sheltzer Lab is supported by NIH grants R01CA237652 and R01CA276666, an American Cancer Society Research Scholar grant, an American Federation for Aging Research New Investigator Award in Aging Biology, and a Colorectal Cancer Alliance Young Investigator award. Research in the Bornhauser Lab is supported by funding source Krebsliga Schweiz KLS-5396-08-2021. Research in the Gao Lab is supported by NIH grant CA225836. E.S. is supported by the National Science Foundation Graduate Research Fellowship. P.L.H. acknowledges support from NIH grant R01GM065313. S.H.N. acknowledges support from NIH grant F32GM150246. S.S. thanks the Humboldt Foundation for a Feodor Lynen Fellowship. Research in the Taylor Lab is funded by Cancer Research UK [DRCPGM\100051] and via funding to the Cancer Research UK Manchester Research Centre through the Cancer Research UK Manchester Centre Award [CANCTA-2023/100004], with additional support from the Medical Research Council [MR/X008088/1 and MR/X502868/1] and the NIHR Manchester Biomedical Research Centre [NIHR203308]; the views expressed are those of the author(s) and not necessarily those of the NIHR or the Department of Health and Social Care. S.R. acknowledges funding from TARGET4OC-GI2-CESPU-2025. S.M.C. acknowledges funding from Bayer and Calico. J.C. is supported by the Howard Hughes Medical Institute.

We thank the Yale Center for Molecular Discovery for their assistance in screening the MDR1-WT and MDR1-KO cells. This work was performed with assistance from the Yale University Flow Cytometry shared resource. These core facilities are supported in part by grant P30CA016359. This research made use of the Chemical and Biophysical Instrumentation Center at Yale University (RRID:SCR_021738). This work was also performed with assistance from the Cold Spring Harbor Laboratory Flow Cytometry shared resource, which is supported in part by grant P30CA045508. We thank Oana C. Danciu (University of Illinois) for sharing the caspase-3 immunohistochemistry data. We thank Paul Hergenrother, Scott Dixon, and members of the Sheltzer Lab for providing helpful feedback on this manuscript.

## Declaration of interests

D.B., K.L., S.N.S., S.S., P.L.H., and J.M.S have filed intellectual property regarding methods to treat drug-resistant cancers based on the results presented in this manuscript. J.M.S. has received consulting fees from Merck, Pfizer, Ono Pharmaceuticals, Highside Capital Management, and Meliora Therapeutics and is a member of the advisory boards of BioIO, Permanence Bio, Karyoverse Therapeutics, and the Chemical Probes Portal. C.R.V. has received consulting fees from Flare Therapeutics, Roivant Sciences, and C4 Therapeutics; has served on the advisory boards of KSQ Therapeutics, Syros Pharmaceuticals, and Treeline Biosciences; has received research funding from Boehringer Ingelheim and Treeline Biosciences; and owns stock in Treeline Biosciences. The other authors declare no competing interests.

## Methods

### Cell lines and culture conditions

The identities of all human cancer cell lines were confirmed by short tandem repeat (STR) profiling performed at the University of Arizona Genetics Core facility. A375, HCT116, MiaPaCa-2, HepG2, A549, and HT29 cells were cultured in Dulbecco’s Modified Eagle Medium (DMEM; Thermo Fisher Scientific, cat. no. 11995-073) supplemented with 10% fetal bovine serum (FBS; Sigma-Aldrich, cat. no. F2442), 2 mM L-glutamine (Lonza, cat. no. 17-605F), and 100 U/mL penicillin-streptomycin (Life Technologies, cat. no. 15140122). Jurkat, Nalm6, LS1034, A2780, H1975, OVK18, Caco-2, OVTOKO, A498, 786O, C4-2B, and OVCAR8 cell lines were maintained in RPMI 1640 medium (Lonza, cat. no. 12-115F/12) supplemented with 10% FBS, 2 mM L-glutamine, and 100 U/mL penicillin-streptomycin. All cell lines were maintained at 37 °C in a humidified atmosphere containing 5% CO₂.

### Generation of knockout clones

To generate CASP3/6/7 triple-knockouts in A375 and HCT116, we followed our established protocol for making knockout clones in cancer cell lines^27^. Briefly, Cas9 was stably expressed via transduction with MSCV_Cas9_puro (Addgene #65655) and selected with puromycin. Vectors targeting either Rosa26 or CASP3 were generated by cloning a gRNA-targeting sequence into LRCherry2.1 (Addgene #108099). A vector targeting both CASP6 and CASP7 was generated using a previously-described system for generating dual-targeting vectors and cloned into Lenti_sgRNA_EFS_GFP (Addgene #65656)^71^. Cells expressing the desired fluorescent marker(s) were single cell sorted onto 96-well plates and expanded before being screened for knockout via western blotting. The gRNA sequences are listed in Table S3A. Jurkat and Nalm6 cells harboring a triple-knockout of caspases 3, 6, and 7 have been previously described^24^. The LS1034 MDR1-KO clones were previously described^29^.

### Western blotting

Cells were harvested, resuspended in RIPA buffer [25 mM Tris, pH 7.4, 150 mM NaCl, 1% Triton X-100, 0.5% sodium deoxycholate, 0.1% sodium dodecyl sulfate, protease inhibitor cocktail (Sigma, cat. no. 4693159001), and phosphatase inhibitor cocktail (Sigma, cat. no. 4906845001)], and vortexed. Protein lysate was quantified via the DC Protein Assay kit (Bio-Rad, cat. no. 5000111). Lysate concentration was equalized, and samples were run on a 10% SDS-PAGE gel at 110 volts. The samples were transferred onto a polyvinylidene difluoride (PVDF) membrane using a Bio-Rad Trans-Blot Turbo Transfer System. The membrane was then blocked for 1 hour at room temperature with 5% milk in PBS. Membranes were incubated overnight at 4°C in primary antibody. The next day, blots were washed 3 times for 10 minutes each in PBST and incubated at room temperature with secondary antibody for one hour. The secondary antibodies used were HRP goat anti-mouse (Bio-Rad, cat. no. 1706516) at a dilution of 1:10,000 and HRP goat anti-rabbit (Abcam, cat. no. ab6721) at a dilution of 1:10,000. Blots were again washed 3 times for 10 minutes in PBST before a 1-minute incubation with ProtoGlow ECL (National Diagnostics, cat. no. CL-300) and developed using autoradiographic film (Lab Scientific, XAR ALF 2025) or imaged electronically (Bio-Rad, ChemiDoc MP). The antibodies used are listed in Table S3B.

### PRISM analysis

Drug sensitivity, gene mutation, gene copy number, and mRNA expression data were downloaded from the DepMap 24Q4 release at www.depmap.org^29,72^. To compute drug similarities, the response of each cell line to each drug was correlated using pairwise complete observations, then Pearson correlations were ranked and plotted in a waterfall plot. Individual drug-drug and drug-gene Pearson correlations were computed using pairwise complete observations of each cell line response to each drug or gene. To compute cell line predictors of PAC-1 sensitivity, Pearson correlations were computed using pairwise complete observations between PAC-1 and (1) each gene mutation for which mutations are present in 2% or more of the drug-treated cell lines, (2) gene copy number, and (3) gene mRNA expression. These correlations were ranked against each other and plotted on a single waterfall plot.

### Cell cycle analysis

About 0.2 million cells were plated onto a 6-well plate. After 24 hours of drug exposure, the cells were harvested by trypsinization followed by centrifugation at 1,000 rpm for 5 min at room temperature. The supernatants were removed and the cell pellets were rinsed with 1X PBS followed by centrifugation at 1,000 rpm for 5 minutes. The supernatants were removed and the cell pellets were rinsed with 1x PBS. Cells were added dropwise to 4 mL of ice cold 100% ethanol and fixed at −20°C for 5 –15 minutes. Fixed cells were pelleted by centrifugation at 1,000 rpm for 5 minutes at room temperature and then washed with 1x PBS. The pellets were then resuspended in PBS containing 0.05% Triton X-100, 10 μg/mL RNase A (Invitrogen, cat. no. A32078), and 20 μg/mL propidium iodide (Life Technologies, cat. no. P3566), filtered through a cell strainer FACS tube cap. Cells were incubated for 30 minutes at room temperature in the dark. Cell cycle analysis was measured using a MACSQuant Analyzer 10 (Miltenyi Biotec, Germany).

### Quantitative real-time PCR

Cells were plated in 6-well plates and treated with the indicated drugs for 24 hours. Total cellular RNA was extracted and isolated using the Qiagen RNeasy Mini Kit (Qiagen, cat. no.74106). cDNA synthesis was performed using SuperScript IV VILO Master Mix (Invitrogen, cat. no. 11756500). PowerTrack™ SYBR Green Master Mix (Thermo Fisher Scientific, cat. no. C46109) was used to perform the qRT-PCR and quantified using the QuantStudio 6 Flex Real-Time PCR system (Applied Biosystems). GAPDH was used for normalization and fold changes were calculated by using the ΔΔC_t_ method. The qPCR primers are listed in Table S3C.

### Quantification of intracellular iron, calcium, and zinc levels

Cells were seeded in 6-well culture plates at appropriate densities and allowed to adhere overnight. Subsequently, cells were treated with the indicated compounds for 24 hours under standard culture conditions. For quantification of total intracellular iron content, we used the iron assay kit (Abcam, cat. no. ab83366) following the manufacturer’s protocol. Briefly, cells were lysed in the provided assay buffer, and iron levels were determined through a colorimetric reaction using a microplate reader. Total intracellular zinc levels were quantified using a zinc assay kit (Cell Biolabs Inc., cat. no. MET-5138). Cells were lysed, and zinc concentrations were determined following the manufacturer’s colorimetric assay protocol. Similarly, intracellular calcium levels were measured using a calcium assay kit (Abcam, cat. no. ab102505), according to the supplier’s instructions. Absorbance readings for these assays were obtained using a microplate spectrophotometer.

### Aconitase (ACO1) activity assay

ACO1 activity was quantified using the Abcam aconitase assay kit (cat. no. ab83459) following the manufacturer’s instructions. Briefly, 1 × 10⁶ A375 cells were treated with the indicated compounds for 24 hours and then washed twice with cold PBS and lysed in assay buffer. The lysate was centrifuged at 800 × g for 10 min at 4 °C, and the supernatant (100 µl) was transferred to a new tube. To activate ACO1, 10 µl of activation solution was added and the mixture was incubated on ice for 1 hour. Activated samples (50 µl per well) were loaded into a 96-well plate, followed by 50 µl of reaction mixture provided in the kit. After incubation at 25 °C for 60 min, 10 µl of developer solution was added and the plate was further incubated at 25 °C for 10 min. Absorbance was measured at 450 nm using a microplate reader.

### Iron and zinc rescue

Cells were plated (5,000 cells/well) in 100 μl media on flat-bottomed 96-well culture plates (CELLTREAT, cat. no. 229196). After 12-24 hours, anti-cancer therapeutics were added to the cells with or without exogenous iron (Millipore Sigma, cat. no. 372870) or zinc (Millipore Sigma, cat. no. Z0251) for 72 to 96 hours. Cell viability was measured by MTS assay (Promega, cat. no. G3580). Data were normalized to the untreated cells and represented by the mean of three independent measurements.

### Drug sensitivity assays

The anti-cancer compounds used in this study are listed in Table S3D. For adherent cancer cell lines, 5,000 cells per well were seeded in 100 μL of complete growth medium into flat-bottomed 96-well culture plates (CELLTREAT, cat. no. 229196). Cells were allowed to adhere for 12–24 hours before drug treatment. Compounds were added starting with the highest concentration in one row, followed by serial dilutions across 7 concentrations. Cell viability was assessed 72–96 hours post-treatment, depending on the proliferation rate of each cell line.

For suspension cell lines, 5,000 cells were seeded per well in 50 μL of medium in round-bottomed 96-well suspension plates (Corning, cat. no. 2797). Cells were treated with anti-cancer drugs diluted in a 50:50 ratio, yielding a final volume of 100 μL per well, with serial dilutions across a 7-point concentration range. Cell viability was measured 96 hours post-treatment.

Cell proliferation and viability were assessed using either a MACSQuant Analyzer 10 flow cytometer (Miltenyi Biotec, Germany) or the MTS assay (Promega, cat. no. G3580), following the manufacturer’s protocols. Viability was normalized to untreated controls and presented as mean values from three to five independent experiments. Only measurements with standard errors below 10% were included in analysis. IC50 values were calculated using a four-parameter inhibition vs. response model in Prism (GraphPad Software, San Diego, CA), based on 7-point dose response curves generated from three to six biological replicates.

### Inductively coupled plasma mass spectrometry analysis

A375 cells (1 × 10⁶) were treated with the indicated drugs for 24 hours, washed twice with 1x PBS, and detached using TrypLE Express Enzyme (Thermo Fisher Scientific). The harvested cells were collected into metal-free 15 mL centrifuge tubes containing an equal volume of complete growth medium to neutralize enzymatic activity. Cells were pelleted by centrifugation, and the supernatant was carefully removed to eliminate residual medium. The cell pellet was subsequently washed once with PBS to remove any remaining culture media components and centrifuged again. For acid digestion, 1 mL of concentrated nitric acid (HNO₃, trace metal grade; Fisher Scientific) was added directly to each pellet. Samples were then incubated at 90°C for 1 hour in a heating block to facilitate digestion, followed by overnight evaporation to dry the samples. The following day, the digested samples were reconstituted to a final concentration of 5% (v/v) HNO₃ with an internal standard indium to ensure analytical accuracy. The samples were subsequently heated at 60°C for 1 hour to ensure complete dissolution. Elemental analysis of metal content was performed using an inductively coupled plasma mass spectrometer (Thermo Fisher Scientific Element XR).

### Lentiviral generation and overexpression in MiaPaCa-2 cells

Expression plasmids encoding ABCB1, ABCC1, and ABCG2 were obtained from VectorBuilder (Chicago, USA). Lentiviral particles were produced by co-transfecting HEK293T cells with 20 µg of the respective expression plasmid, 25 µg of the packaging plasmid psPAX2, and 4 µg of the envelope plasmid pVSVG in a 100 mm culture dish seeded with approximately 3 million HEK293T cells. At 48 hours post-transfection, the virus-containing supernatant was collected and passed through a 0.45 µm sterile syringe filter to remove cellular debris. The filtered lentiviral supernatant was then used to transduce MiaPaCa-2 cells in the presence of 8 µg/mL polybrene (Santa Cruz Biotechnology, cat. no. SC-134220) to enhance infection efficiency. After a 24-hour incubation period, the transduction medium was replaced with fresh complete growth medium. Successful overexpression of ABCB1, ABCC1, and ABCG2 was confirmed by qRT-PCR.

### Analysis of patient-derived ovarian cancer cell models

Research samples were obtained from the Manchester Cancer Research Centre (MCRC) Biobank, UK. The MCRC Biobank holds a generic ethics approval which can confer this approval to users of banked samples via the MCRC Biobank Access Policy (Human Tissue Authority license: 30004; South Manchester Research Ethics Committee ref: 22/NW/0237). Further details regarding ethical governance and licensing can be found at: https://www.mcrc.manchester.ac.uk/research/mcrc-biobank/accessing-the-mcrc-biobank/. Ovarian Cancer Models (OCMs) were established as described previously^42^. Briefly, ascites were centrifuged, red blood cells removed (Red Blood Cell Lysis Solution, Miltenyi Biotec) and remaining cells plated into Primaria or Cell+ flasks containing OCMI^73^. Cultures were incubated at 37°C for 2–4 days in a humidified 5% CO2 and 5% O2 atmosphere, then media replaced every 3–4 days. Once attached, selective trypsinisation was used to separate stromal and tumour cells. Established OCMs were cultured in OCMI^42,73^.

Drugs were made up in DMSO (Sigma) with stocks aliquoted and stored at -80 °C. Cells were seeded at 2000 cells/well into 12-well Cell+ plates (Sarstedt). The following day a titration of PAC-1 (1, 2.5, 5, 7.5, 10, 15, 20, 25 µM, SelleckChem) was added and cells incubated with the drug continually for 21 days, with the media changed every 5–7 days. Plates were fixed in 1% formaldehyde, before being stained with a 0.05% v/v crystal violet (Sigma Aldrich) solution. Stained plates were imaged on a ChemiDoc™ Touch Imaging System (BioRad) before the crystal violet was extracted with 10% acetic acid. Absorbance at 570 nm was read on a VarioSkan™LUX multimode microplate reader (Thermo Scientific). The mean absorbance from six 10% acetic acid only blank wells was subtracted from each value, before normalization to the untreated control well. Normalized values from at least three biological replicates were plotted against Log10 PAC-1 concentration.

RNA sequencing of OCMs was used to measure ABCB1 expression levels^43^. Total RNA extracted using a RNeasy Plus Mini kit (Qiagen) was submitted to the Genomic Technologies Core Facility at The University of Manchester. After quality and integrity assessment using a 4200 TapeStation (Agilent Technologies), libraries were generated using the Illumina® Stranded mRNA Prep Ligation kit (Illumina, Inc.) according to the manufacturer’s protocol. Briefly, polyadenylated mRNA was purified from 0.025¬–1 µg total RNA using poly-T oligo-attached magnetic beads. mRNA was fragmented at elevated temperature before reverse transcription into first-strand cDNA using random hexamer primers in the presence of Actinomycin D. Following removal of template RNA, second-strand cDNA was synthesised to yield blunt-ended, double-stranded cDNA fragments (with strand specificity maintained by dUTP incorporation in place of dTTP to quench the second strand during subsequent amplification). Following a single adenine base addition, adapters with a complementary thymine overhang were ligated to the cDNA fragments followed by ligation of pre-index anchors to prepare for dual indexing. Index adapter sequences were added by PCR to enable multiplexing of the final cDNA libraries, which were pooled and loaded onto an SP flow cell and paired-end sequenced (59 + 59 cycles, plus indices) on an Illumina NovaSeq6000 instrument. Output data were demultiplexed and binary base call (BCL)-to-Fastq conversion performed using bcl2fastq software (Illumina, Inc., v2.20.0.422). Stranded paired-end reads were quality assessed using FastQC (v0.11.3)^74^ and FastQ Screen (v0.14.0)^75^, followed by adapter and low-quality base trimming with BBDuk from the BBMap suite (v36.32)^76^. Trimmed reads were mapped against the human reference genome (hg38) and gene annotation from Gencode (v32) using STAR (v2.7.2b)^77^. The “–quantMode GeneCounts” option was used to obtain read counts per gene from STAR. The R package DESeq2^78^ was then used to apply the median of ratios method of normalization.

### Generation of SLC25A40-ABCB1 fusion cell line models

gRNAs targeting the regions of SLC25A40 and ABCB1 fusion breakpoints identified in a patient-derived tumor^14^ were designed using Benchling. A gBlock gene fragment (Integrated DNA Technologies) containing the two gRNAs was cloned into a Lenti-Cas9-gRNA-GFP vector (Addgene #124770). gRNA sequences and chromosome positions are listed in Table S3A.

A2780 cells were transiently transfected with the constructs using Lipofectamine® 3000 (Thermo Fisher Scientific, cat. no. L3000015). After 48 hours, GFP-positive single cells were isolated using a BD FACSAria™ III Cell Sorter and expanded for approximately two weeks. Genomic DNA was extracted using the DNeasy® Blood & Tissue Kit (Qiagen, cat. no. 69506), and fusion junctions were verified by PCR with primers flanking the breakpoints. PCR products were separated on a 0.8% agarose gel and visualized with the Bio-Rad GelDoc Go Imaging System (Bio-Rad, cat. no. 12009077). Bands of the expected size (∼500–700 bp) were excised and purified using the GeneJET Gel Extraction Kit (Thermo Scientific, cat. no. K0692). Purified products underwent linear/PCR sequencing by Plasmidsaurus, followed by custom analysis and annotation. MDR1 expression in the resulting clones was quantified by qPCR and rhodamine efflux assay.

### Rhodamine efflux assay

Wild-type and MDR1 knockout LS1034 cells were cultured in 6-well plates and loaded with 2.5 μM rhodamine-123 (Thermo Fisher Scientific, cat. no. R302) for 20 minutes at 37°C in 5% CO2. After the accumulation period, the cells were washed three times with PBS. Efflux was initiated by resuspending the cells in rhodamine-free medium with or without the indicated compounds. Following a 60-minute incubation at 37°C, the cells were washed twice with PBS. Rhodamine efflux was quantified using a MACSQuant Analyzer 10 (Miltenyi Biotec, Germany). The rhodamine-123 efflux assays were performed using three independent biological replicates, in which each biological replicate had two technical replicates per condition.

### Competition assay

LS1034 wild-type and MDR1-knockout cell lines were transduced with blue fluorescent protein (BFP) and red fluorescent protein (RFP) expression plasmids, respectively. Following transfection, cells were subjected to bulk fluorescence-activated cell sorting to enrich populations based on BFP and RFP expression, ensuring distinct fluorescently labeled populations.

For competition assays, equal numbers of BFP-positive (BFP⁺) and RFP-positive (RFP⁺) cells were co-seeded, with 25,000 cells of each population plated together into individual wells of a 6-well plate. After 16 hours, the mixed cell populations were treated with the specified drugs at the indicated concentrations. After 72 hours post treatment, cells were harvested and analyzed using a MACSQuant Analyzer 10 flow cytometer (Miltenyi Biotec, Germany). The proportions of BFP⁺ and RFP⁺ cells within the mixed populations were quantified based on fluorescence intensity. Data were represented as the percentage of BFP-positive and RFP-positive cells relative to the total number of fluorescent cells.

### Characterization of chemotherapy-resistant cell lines

Docetaxel-resistant C4-2B cells, characterized by high expression levels of ABCB1 gene, were obtained as a kind gift from Dr. Allen C. Gao’s laboratory at the University of California, Davis^39^. Paclitaxel-resistant OVCAR8 cells (OVCAR8-PTX) were generously provided by Dr. Sara Ricardo’s laboratory at the University of Porto, Portugal^53^. ABCB1 overexpression and paclitaxel resistance in the Docetaxel-resistant C4-2B cells and Paclitaxel-resistant OVCAR8-PTX cells were confirmed in our experimental setting, ensuring the suitability of these cell models for subsequent drug resistance and efflux studies.

### Evolution assay

To investigate the adaptive regulation of ABCB1 expression upon prolonged drug exposure, an evolution assay was conducted using ABCB1-overexpressing OVCAR8 (ovarian cancer) cell lines, as well as the Caco-2 (colon cancer) cell line, which inherently exhibits high endogenous ABCB1 expression. Cells were continuously cultured in the presence of a sub-lethal dose of PAC-1, a concentration empirically determined to exert partial growth inhibition without inducing significant cytotoxicity. The treatment was maintained for a total of 12-15 passages, corresponding to an approximate duration of 50-60 days. At every third passage, a fraction of the cell population was collected for RNA extraction to monitor the temporal changes in ABCB1 mRNA expression levels. Upon completion of the long-term evolution assay (approximately 50 days for OVCAR8-PTX, spanning 12 passages, and 60 days for Caco-2, spanning 15 passages), the derived cell populations were subjected to paclitaxel sensitivity assays to assess changes in drug responsiveness induced by prolonged PAC-1 exposure.

### Efflux assay

The efflux assay was performed by Eurofins, USA, using hMDR1-MDCKII_cMDR1_KO cells, a Madin-Darby canine kidney cell line stably expressing the human MDR1 gene. Cells were seeded at a density of 5.24 × 10^5^ cells/cm² (equivalent to 1.0 × 10^6^ cells/mL; 75 μL/well) onto porous polycarbonate membranes in 96-well Corning HTS Transwell® plates. Monolayers were used 3–5 days post-seeding. This assay was performed in the apical to basolateral (A-B) and the basolateral to apical (B-A) directions. The test compounds were prepared at 10 μM in FaSSIF (fasted state simulated intestinal fluid) pH 6.5 on the apical side and HBSS-HEPES with 0.5% dialyzed FBS (pH 7.4) on the basolateral side and final DMSO concentration of 1%. Test compounds were prepped in either the presence or absence of 100 μM FeCl_2_ to be added on both sides of monolayer.

Prior to the assay, cells were washed and pre-incubated with HBSS buffer to remove residual media components. Test solutions were equilibrated by shaking and warming before being applied to the assay plates. Plates were incubated at 37 °C with gentle shaking for 120 min. Samples were collected from the donor compartment at t = 0 and t = 120 min, and from the receiver compartment at t = 120 min. Samples were precipitated in acetonitrile with 0.1% formic acid. Stop plates were centrifuged and supernatant sampled for LCMS analysis. Peak areas are obtained for calculations using the equation below. Known P-gp substrate Digoxin was used as a control. Cells are incubated post assay with zero marker fluorescein to ensure monolayer integrity, wells failing FL tests are removed from data sets.

The apparent permeability coefficient (P_app_) of the test compound and its recovery are calculated as follows:

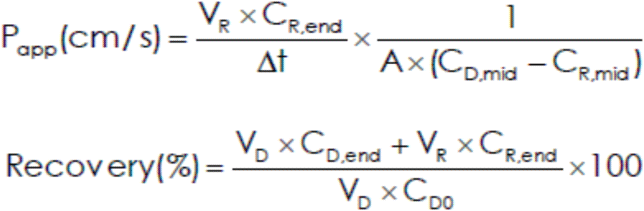

A is the surface area of the cell monolayer (0.11 cm2).

C is concentration of the test compound, expressed as peak area.

D denotes donor and R is receiver.

0, mid, and end denote time zero, mid-point, and end of the incubation.

Delta t is the incubation time.

V is the volume of the donor or receiver.

For the transwell efflux assay, two biological replicates with two technical replicates for each condition were performed. The results represent the mean of the independent biological assays.

### High-throughput drug screen and analysis

For the high-throughput screen to identify MDR1-targeting cancer drugs, 1,000 cells/well were plated on black, clear-bottom 384-well plates (Corning cat. no. 3764) using MultiDrop Combi and allowed to adhere overnight. The next day, an Echo acoustic dispenser (Beckman) was used to add compounds in 8 serial dilutions with 10 μM as a top concentration. The toxicity of 129 anti-cancer compounds (NCI Oncology Set 5 library), PAC-1, and paclitaxel, were assessed. After 72 hours of drug exposure, cell viability was assessed using CellTiter Glo. Test compound data were normalized relative to a 0.1% DMSO negative control and a 60 μM Bortezomib positive control. The IC50 values were generated via the high-throughput screen or from confirmatory MTS assays. A waterfall plot was generated from (1) log2 transforming the IC50s of LS1034 wild-type, MDR1-KO clone 6, and MDR1-KO clone 9, (2) dividing the transformed WT IC50 by the average of the transformed MDR1-KO clones, and (3) plotting the rank of IC50 deltas.

### Protein expression and purification

The synthetic full-length human *ABCB1* gene (residues 1-1280) was codon-optimized for mammalian cell expression. The wild type protein was cloned into a BacMam expression vector containing a C-terminal PreScission cleavage site followed by a green fluorescence protein (GFP) tag. For protein expression, baculovirus carrying recombinant ABCB1 construct was amplified in Sf9 cells; HEK293F cells were infected with recombinant baculovirus, grown at 37 °C for 8 h, induced with 10 mM sodium butyrate, and then were grown at 30 °C for 48 h before harvesting. Cell pellets were resuspended by manual homogenization with a dounce in lysis buffer containing 50 mM HEPES pH 8.0, 150 mM KCl, 20% glycerol, 5 mM MgCl2, 1 mM DTT, 1 mM Mg^2+^/ATP, and protease inhibitors (1 mM PMSF, 1 mM benzamidine, 0.1 μg/mL trypsin inhibitor, 0.1 μg/mL pepstatin A, 0.1 μg/mL leupeptin, 0.1 μg/mL aprotinin, and 3 μg/mL DNase). Membranes were solubilized by adding 1% n-dodecyl-β-D-maltopyranoside (DDM) and incubating at 4 °C for 75 min. Solubilized membranes were centrifuged at 75,000 x g for 45 min to remove the insoluble fraction, and the supernatant was loaded onto pre-equilibrated NHS-activated Sepharose 4 Fast Flow bead (GE Healthcare) coupled to an anti-GFP nanobody. The resin was washed with 20 column volumes of 50 mM HEPES pH 8.0, 150 mM KCl, 0.05% DDM, 0.01% cholesteryl hemisuccinate (CHS), 1 mM DTT, and 1 mM Mg^2+^/ATP and then incubated with PreScission protease (10:1 w/w ratio) at 4 °C for 2 hr. The eluted protein was reloaded onto an anti-GST column to remove the protease. Eluate was concentrated using an Amicon Ultra (MWCO 100K, Millipore) centrifugal device and then purified by Superose 6 Increase 10/300 column (GE Healthcare) in buffer containing 50 mM HEPES pH 8.0, 150 mM KCl, 0.05% DDM, 0.01% CHS, and 1 mM DTT.

### ATP hydrolysis measurements

ATP hydrolysis by MDR1 was measured at 37 °C by quantifying the release of inorganic phosphate as described^79^. The reaction mixture consisted of 2 μg detergent purified MDR1, 50 mM HEPES pH 8.0, 150 mM KCl, 0.05% DDM, 0.01% CHS, 2 mM MgCl2. For substrate-stimulated conditions, the solution was incubated with substrate on ice for 30 min, and subsequently prewarmed for 5 min at 37°C. ATP was added to the mixture at a final concentration of 4 mM, and the reaction was initiated in a final volume of 50 uL at 37°C. The reaction was stopped after 20 min by addition of 50 μL of 12% SDS, and the color was initially developed by adding 100 μL of 1% ammonium molybdate and 6% ascorbic acid in 1 M HCl and incubating for 5 minutes at room temperature. The color was further developed by addition of 150 μl solution containing 2% (w/v) sodium citrate, 2% (w/v) sodium metarsenite, and 2% (v/v) acetic acid and incubation for 10 min at 37°C. The absorbance at 850 nm was measured using an Infinite M1000 microplate reader (Tecan). The amount of hydrolyzed phosphate was determined by comparison to standard curve of potassium phosphate (KH_2_PO_4_). Data were fitted into Michaelis-Menten equations using the GraphPad Prism 10 software.

### PAC-1: iron structure characterization

#### Overview

Spectroscopic chelation assays have shown that PAC-1 and some analogues can bind to Fe^2+^, but the stoichiometry and binding mode have not been determined^35^. To identify the binding mode, we first studied the chelation of Fe^2+^ with PAC-1 in 50 mM HEPES buffer at pH 7.4. Mixtures gave bands between 300 and 500 nm, indicating coordination (Figure 3 and S4B). We used the method of continuous variation with UV-vis spectroscopy (Figure 3B) to determine that the binding stoichiometry was 1:1 between PAC-1 and Fe^2+^, which is also consistent with ^1^H NMR and Mössbauer spectra (Figure S4C-E). This is the same as the reported binding stoichiometry between PAC-1 and Zn^2+25^.

The structure of a 1:1 complex was elucidated by X-ray crystallography (Figure 3A and S4A). Treatment of two equivalents of PAC-1 with FeCl_2_ in a methanol/ethanol mixture gave a red crystalline solid in 85% yield (relative to Fe). The crystallographic structure reveals a remarkable decamer of PAC-1 and Fe^2+^ in a 1:1 ratio; though the solution structure may differ, the stoichiometry agrees with the spectroscopic assessment above. In the crystal structure, electron density adjacent to the N–benzyl nitrogen of each piperazine suggests that the piperazine moieties are protonated, so it acted as a Brønsted base toward another molecule of PAC-1, which is deprotonated at the phenol group. Chloride ions are also present, indicating protonation of basic functional groups. The solid-state ^57^Fe Mössbauer spectrum (Figure S4D-E) of the crushed crystals has two components in an approximately 7:3 ratio (δ_1_ = 1.04 mm/s; Δ*E*Q1 = 2.79 mm/s; δ_2_ = 1.08 mm/s; Δ*E*Q2 = 3.10 mm/s). The two components are consistent with high-spin Fe^2+^ centers in slightly different local coordination environments, which is supported by the crystallographic data (see below for discussion).

While the decameric structure that we characterized in the solid state may break up in solution, it shows clearly that PAC-1 can coordinate to Fe^2+^ through the phenol O, the amide O, and the hydrazide N. To test whether this binding mode is also present in solution, we compared the UV-vis spectra of Fe^2+^ with PAC-1 to two structurally related analogues PAC-2 (which lacks the allyl group) and PAC-1a (which lacks the allyl and hydroxyl groups) at pH 7.4. PAC-2 chelates Fe^2+^ like PAC-1, as indicated by the broad bands between 300 and 500 nm. In contrast, PAC-1a, in which the hydroxyl group is absent, shows no spectroscopic changes that indicate binding. This suggests that the hydroxyl group enables tridentate iron chelation. This is consistent with previous work with Zn^2+^, where the hydroxyl group was necessary for coordination^25,80^.

The crystal structure also shows that the piperazine moiety of PAC-1 can coordinate iron in a bidentate mode (N,N via the hydrazide and piperazine). By analogy, it is possible that the piperazine moiety enables an additional chelation site for other metal ions as well (e.g. Zn^2+^, Na^+^, K^+^). The crystal structure also shows that the nitrogen(s) of the piperazine can act as Brønsted bases, and may be protonated under certain biological conditions. Previous structure–activity relationship studies demonstrated the dependence of anti-tumor activity on both the presence of the piperazine moiety and the substituents bound to it^20^. In this context, our results show that the piperazine moiety can bind to both protons and metal centers, and thus variations that influence the p*K*_a_ and Lewis basicity are expected to have significant impacts on the coordination chemistry and overall activity *in vivo* as well.

#### Methods for synthetic iron(II) compounds

The synthetic iron(II)-PAC-1 compounds were extremely air-sensitive, and all manipulations of these compounds were carried out under an N_2_ atmosphere using either standard Schlenk techniques or a nitrogen-filled glovebox (Vigor Tech). All solvents and reagents were stored inside the glovebox. Toluene was purified by passage through activated alumina columns under an argon atmosphere and stored over activated 4 Å molecular sieves. Dimethyl sulfoxide-*d*_6_ (DMSO-*d*_6_) was purchased from Cambridge Isotope Laboratories, degassed by sparging with nitrogen, and stored over activated 4 Å molecular sieves inside the glovebox. Methanol was sparged with N_2_ and dried by storage over 3 Å molecular sieves inside the glovebox for at least 72 h. Buffer A (50 mM HEPES, 100 mM KNO_3_, pH = 7.4) was prepared from HEPES buffer solution (pH 7.4, Boston BioProducts, Inc.) and KNO_3_ (Sigma-Aldrich) and sparged with N_2_ before use. FeCl_2_ • 4H_2_O was purchased from Strem chemicals and used without further purification. ^1^H NMR data were collected on an Agilent 500 MHz spectrometer. Chemical shifts in ^1^H NMR spectra are referenced to the residual protiated solvent peaks of DMSO-*d*_5_ (2.50 ppm). Mössbauer samples were placed in Delrin sample cups and recorded on a SEECo Mössbauer spectrometer with alternating constant acceleration; isomer shifts are relative to α-iron metal at 298 K. The sample temperature was maintained at 80 K in a Janis Research Company Inc. cryostat. The zero-field spectra were simulated using Lorentzian doublets using MossA. UV/Vis spectra were collected on an Agilent Cary 60 or an Agilent Cary 5000 spectrophotometer.

#### UV-vis measurement of binding of different structural analogues

Aliquots from a 1 mM FeCl_2_ stock solution in buffer A were diluted into 2 mL of buffer A to give final concentrations of 50 µM. Aliquots of PAC-1, PAC-1a or PAC-2 from 10 mM DMSO stock solutions were added to give final concentrations of 50 µM. After 5 minutes, the absorbance spectra were measured in air-tight quartz cuvettes with a path-length of 10 mm.

#### Method of continuous variation

A series of solutions of PAC-1 (from DMSO stock solution) and FeCl_2_ were prepared in buffer A such that the sum of metal and PAC-1 concentrations was kept constant at 50 μM. The mole fraction of PAC-1 was varied between 0.1 and 1.0. After 5 minutes of mixing, absorbance spectra were measured in quartz cuvettes of 10 mm path length. Different backgrounds were collected to account for the different concentrations of DMSO. A 0.2 Hz low-pass FFT filter as implemented in Origin 2020^TM^ was applied to the data to reduce noise. The absorbance at 386 nm was plotted against the mole fraction of PAC-1 (Figure S4B).

#### Synthesis of [(PAC-1)Fe^II^(ROH) • HCl]_10_

In the glove box, FeCl_2_ • 4H_2_O (14.9 mg, 0.075 mmol, 1.0 equiv) was weighed into a 1 dram vial and was dissolved in MeOH (1.0 mL). PAC-1 (60.3 mg, 0.158 mmol, 2.1 equiv) was weighed into a separate 1 dram vial, and the FeCl_2_ solution was added in one portion to the solid PAC-1 using a glass pipette; the suspension immediately became dark red in color. After stirring vigorously for 1 minute, all the PAC-1 had dissolved, and the solution became homogeneous. The solution was then passed through an oven-dried glass fiber filter into a separate, tared 1 dram vial. The vial was then placed inside a –10 °C freezer in the glove box for 16 hours, after which time, small, dark red crystals had formed. The colorless mother liquor was decanted away, and the crystals were washed with 0.25 mL of cold MeOH. After drying under vacuum, the complex was isolated as a dark red crystalline solid (28.1 mg, 0.054 mmol (5148.25 g/mol for Fe_10_C_240_H_310_N_40_O_30_Cl_10_, assuming ROH = MeOH), 72% yield).

#### ^1^H NMR (500 MHz, DMSO-*d*_6_)

δ 90.17 (s, 1H), 67.52 (s, 1H), 50.77 (s, 1H), 15.42 (s, 2H), 7.61 – 7.36 (m, 1H), 7.27 (d, *J* = 27.6 Hz, 3H), 7.04 (s, 3H), 6.71 (s, 2H), 6.05 – 5.81 (m, 1H), 5.15 – 4.96 (m, 1H), 4.09 (d, *J* = 6.0 Hz, 1H), 3.14 – 3.05 (m, 2H), 0.42 (s, 4H), -2.67 (s, 4H), -4.97 (s, 1H), -10.82 (s, 1H). Shown in Figure S4C.

#### Mössbauer spectroscopy

The Mössbauer spectrum of solid [(PAC-1)Fe^II^(ROH) • HCl]_10_ could be fit in two ways, each of which has two components. These two components have only slight differences δ and |Δ*E*_Q_| values. All fitted components are consistent with high spin Fe^2+^. The presence of slightly different environments is consistent with the crystal structure of the [(PAC-1)Fe^II^(ROH) • HCl]_10_ complex, as 1) the identity of the ROH ligand can differ and 2) it is possible that the alcohol coordinated to the Fe center may be deprotonated in certain cases. A more detailed discussion of the uncertainty and disorder in the crystallographic data is discussed below.

#### Mössbauer (80 K, solid, fit 1)

δ_1_ = 1.04 mm s^−1^, |Δ*E*_Q1_| = 2.79 mm s^−1^, int_1_ = 75%; δ_2_ = 1.07 mm s^−1^, |Δ*E*_Q2_| = 3.10 mm s^−1^, int_2_ = 25%. Shown in Figure S4D.

#### Mössbauer (80 K, solid, fit 2)

δ_1_ = 0.96 mm s^−1^, |Δ*E*_Q1_| = 2.85 mm s^−1^, int_1_ = 35%; δ_2_ = 1.11 mm s^−1^, |Δ*E*_Q2_| = 2.91 mm s^−1^, int_2_ = 65%. Shown in Figure S4E.

#### X-ray crystallography

In the glovebox, FeCl_2_ • 4H_2_O (4.9 mg, 0.025 mmol, 1.0 equiv) was weighed into a 1 dram vial and was dissolved in a mixture of MeOH (0.75 mL) and EtOH (0.25 mL). PAC-1 (20.0 mg, 0.051 mmol, 2.05 equiv) was weighed into a separate 1 dram vial and was dissolved in toluene (0.25 mL). The FeCl_2_ solution was added in one portion to the PAC-1 solution using a glass pipette; the solution immediately became dark red in color. After stirring vigorously for 1 minute, the solution was filtered through an oven-dried glass fiber filter into a separate 1 dram vial. The solution was placed inside a –10 °C freezer for 48 hours, after which time large red crystals had formed along the walls of the vial. The mother liquor was removed and the crystals were suspended in Krytox oil. The vial was sealed, removed from the glovebox, and kept cold on dry ice while transporting to the diffractometer.

Low-temperature diffraction data (ω-scans) were collected on a Rigaku Synergy-S diffractometer coupled to a HyPix-Arc 100 detector with Cu Kα (λ = 1.54178 Å) for the structure of syn-24076. The diffraction images were processed and scaled using Rigaku Oxford Diffraction software (CrysAlisPro; Rigaku OD: The Woodlands, TX, 2015). The structure was solved with SHELXT and was refined against *F*^2^ on all data by full-matrix least squares with SHELXL^81^. All non-hydrogen atoms were refined anisotropically. Hydrogen atoms were included in the model at geometrically calculated positions and refined using a riding model. The isotropic displacement parameters of all hydrogen atoms were fixed to 1.2 times the U value of the atoms to which they are linked (1.5 times for methyl groups).

With chlorides present in the structure, both the alcohol and piperazines would be protonated to balance the overall charge. Residual electron density adjacent to the N–benzyl nitrogens of the piperazine moieties further supports protonation of these sites. The protons associated with the piperazines and alcohols coordination to the iron centers were placed in geometrically expected positions based on chemical expectations. Other structural models are possible, but given the large solvent voids and diffuse difference map, this model fits best with other supporting evidence.

There are numerous sites of disorder within this model. In general, the following approach was used to create stable models for refinement. The site occupancies of the disordered atoms were freely refined to converged values. In all cases, the population distribution of the models ranged from 0.50/0.50 to 0.60/0.40. In the case of one benzylamine, the site occupancies were constrained to be exactly 0.50/0.50. Chemically similar, disordered C-C, C-N and C-O bonds were restrained to have similar distances. The thermal parameters of disordered groups were restrained to have similar displacement values. The standard uncertainties (SU) of most restraints were set to 0.02 (i.e. the default SHELXL value). In some cases, the SU was set to 0.002 to achieve a stable refinement. There appears to be a distribution of 6:4 ethanol:methanol solvents coordinated to iron in the asymmetric unit. However, these assignments are tenuous as the libration and disorder obscures the true identity and distribution of coordinating solvents.

The interstitial spaces between and within the macrocycles create large voids that are occupied by a mixture of ethanol and methanol. The total void space was calculated to be 36.2% (6760 Å^3^) of the unit cell^82^. Many sites were found and their initial positions within the asymmetric unit are copied below.

The most likely chloride sites were found. One appeared to be disordered and accounted for 0.62 of a chloride. The next largest difference map peak was set as its counterpart with a linked free variable. The remaining difference map density was removed in preparation for solvent masking.

A program based on BYPASS^83^ was used to compensate for the contribution of disordered solvents contained in voids within the crystal lattice from the diffraction intensities. This procedure was applied to the data file and the submitted model is based on the solvent-removed data. Based on the total electron density found in the voids (916 e/Å^3^), we estimate that 16 ethanol and 16 methanol molecules are present in the unit cell.

Several reflections were found to have a high error/standard deviation and were omitted from the least square refinement. Some reflections were very close to one another and streaky which complicated the integration. The crystal also had some small twin components that were ignored. These factors, possibly among others, contributed to the large number of bad reflections.

*Possible solvent positions (either full or partially occupied):*

O1x 6 0.33946 1.09999 0.83899 11.00000 0.11852

C1x 1 0.39836 1.08461 0.81615 11.00000 0.16627

C2x 1 0.41082 1.12817 0.79398 11.00000 0.19306

O2x 6 0.65455 1.32880 0.73228 11.00000 0.07258

C3x 1 0.64557 1.31584 0.77199 11.00000 0.17951

C4x 1 0.72638 1.32025 0.78763 11.00000 0.24746

O3x 6 0.82137 1.28311 0.89175 11.00000 0.17238

O4x 6 0.88714 1.17237 0.98021 11.00000 0.12389

O5x 6 0.71466 0.81179 1.13551 11.00000 0.06366

C5x 1 0.66930 0.77078 1.11399 11.00000 0.07318

O6x 6 0.40556 0.74073 0.76882 11.00000 0.10301

C6x 1 0.41660 0.76843 0.81317 11.00000 0.13650

C7x 1 0.40378 0.82276 0.81535 11.00000 0.21033

O7x 6 0.46147 0.76628 0.70166 11.00000 0.10273

C8x 1 0.52560 0.79049 0.71473 11.00000 0.10429

O9x 6 0.64020 0.83497 0.43797 11.00000 0.11274

C9x 1 0.65008 0.88638 0.43410 11.00000 0.13580

C10x 1 0.62557 0.91931 0.46248 11.00000 0.14703

o10x 6 0.89859 1.24690 0.83329 11.00000 0.06792

C26x 1 0.92650 1.29011 0.81900 11.00000 0.09914

O11x 6 0.54056 0.71966 0.57952 11.00000 0.14941

C11x 1 0.55299 0.66755 0.59482 11.00000 0.15547

O12x 6 0.52526 0.64600 0.76441 11.00000 0.10377

C12x 1 0.52850 0.65542 0.72360 11.00000 0.14583

C13x 1 0.52742 0.61105 0.69362 11.00000 0.14650

O13x 6 0.57615 0.56592 0.78797 11.00000 0.08128

C14x 1 0.63742 0.57685 0.81579 11.00000 0.16736

C15x 1 0.65083 0.53102 0.83914 11.00000 0.16736

O14x 6 0.65047 0.43761 0.84417 11.00000 0.40719

C16x 1 0.67127 0.43756 0.89441 11.00000 0.31187

C17x 1 0.69301 0.48307 0.93195 11.00000 0.39785

O15x 6 0.94439 0.61164 0.97615 11.00000 0.12412

C18x 1 0.89945 0.59262 0.99713 11.00000 0.14296

C19x 1 0.87492 0.54288 0.97767 11.00000 0.22178

O18x 6 1.01154 0.93480 0.64422 11.00000 0.20905

C25x 1 0.96193 0.89824 0.60458 11.00000 0.22768

O19x 6 0.87088 0.44995 0.52490 11.00000 0.07406

C23x 1 0.92674 0.44601 0.55005 11.00000 0.12516

C24x 1 0.97381 0.48454 0.55523 11.00000 0.13099

O20x 6 0.83129 0.47209 0.67292 11.00000 0.17089

C22x 1 0.85133 0.50825 0.70999 11.00000 0.14270

C20x 1 0.96889 0.60797 0.79955 11.00000 0.18938

O21x 6 0.97637 0.65754 0.83229 11.00000 0.19335

C21x 1 1.03046 0.58991 0.79688 11.00000 0.40458

O22x 6 1.08148 0.87429 0.79619 11.00000 0.11789

C27x 1 1.14500 0.88575 0.78969 11.00000 0.12220

C28x 1 1.19128 0.86501 0.82312 11.00000 0.20347

The full numbering scheme of compound syn-24076 can be found in the full details of the X-ray structure determination (CIF), which is included as Supporting Information. CCDC number 2359795 (syn-24076) contains the supplementary crystallographic data for this paper. These data can be obtained free of charge from The Cambridge Crystallographic Data Center via www.ccdc.cam.ac.uk/data_request/cif.

## Supplemental Figure Legends

**Figure S1.**
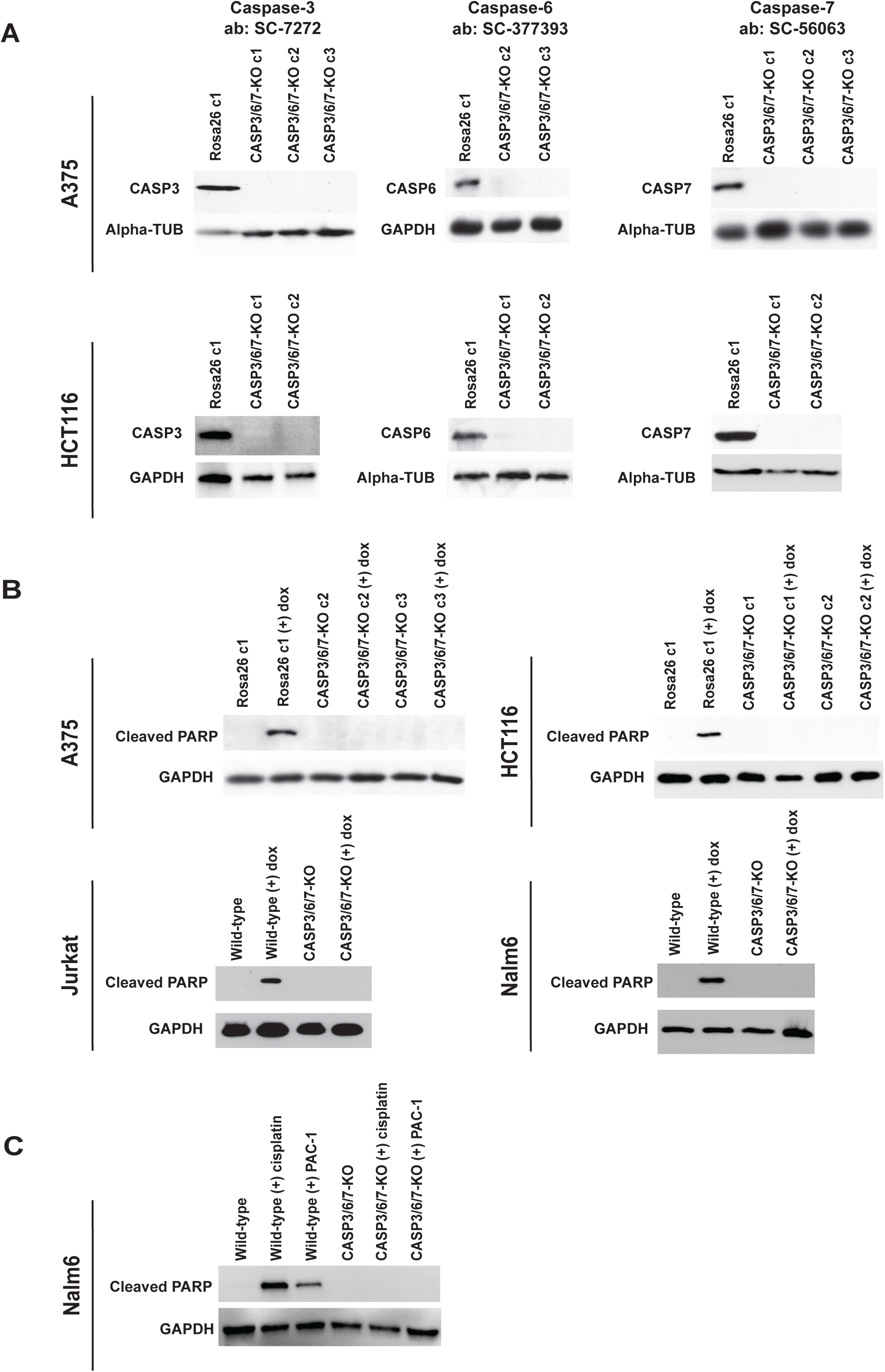
Validation of executioner caspase ablation. (A) Western blot validation of the loss of caspase-3, caspase-6, and caspase-7 expression in the indicated cell lines using an additional set of antibodies. Alpha-tubulin or GAPDH levels were examined as a loading control. (B) Western blot analysis of PARP cleavage in cells treated with 2 μM doxorubicin. GAPDH levels were examined as a loading control. (C) Western blot analysis of PARP cleavage in cells treated with 5 μM cisplatin or 10 μM PAC-1. GAPDH levels were examined as a loading control.

**Figure S2.**
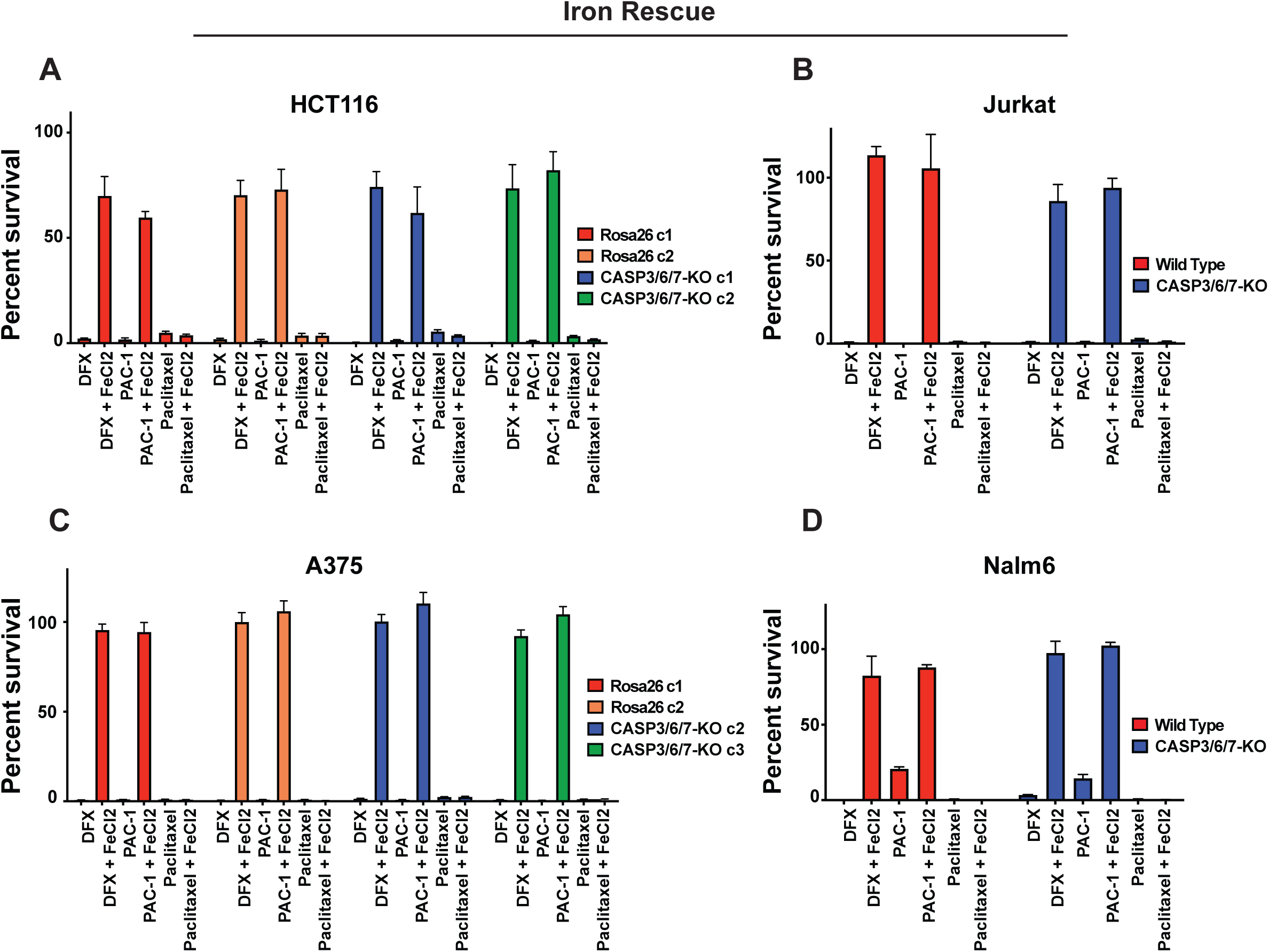
Iron supplementation restores cellular viability in wild-type and CASP3/6/7-KO cells treated with PAC-1. (A) Cellular viability in HCT116 wild-type and CASP3/6/7-KO clones treated with 10 μM deferasirox (DFX), 10 μM PAC-1, or 20 nM paclitaxel, in the presence or absence of 150 µM FeCl_2_. (B) Cellular viability in Jurkat wild-type and CASP3/6/7-KO clones treated with 10 μM deferasirox (DFX), 5 μM PAC-1, or 5 nM paclitaxel, in the presence or absence of 150 µM FeCl_2_. (C) Cellular viability in A375 wild-type and CASP3/6/7-KO clones treated with 10 μM deferasirox (DFX), 10 μM PAC-1, or 30 nM paclitaxel, in the presence or absence of 150 µM FeCl_2_. (D) Cellular viability in Nalm6 wild-type and CASP3/6/7-KO clones treated with 10 μM deferasirox (DFX), 5 μM PAC-1, or 5 nM paclitaxel, in the presence or absence of 150 µM FeCl_2_.

**Figure S3.**
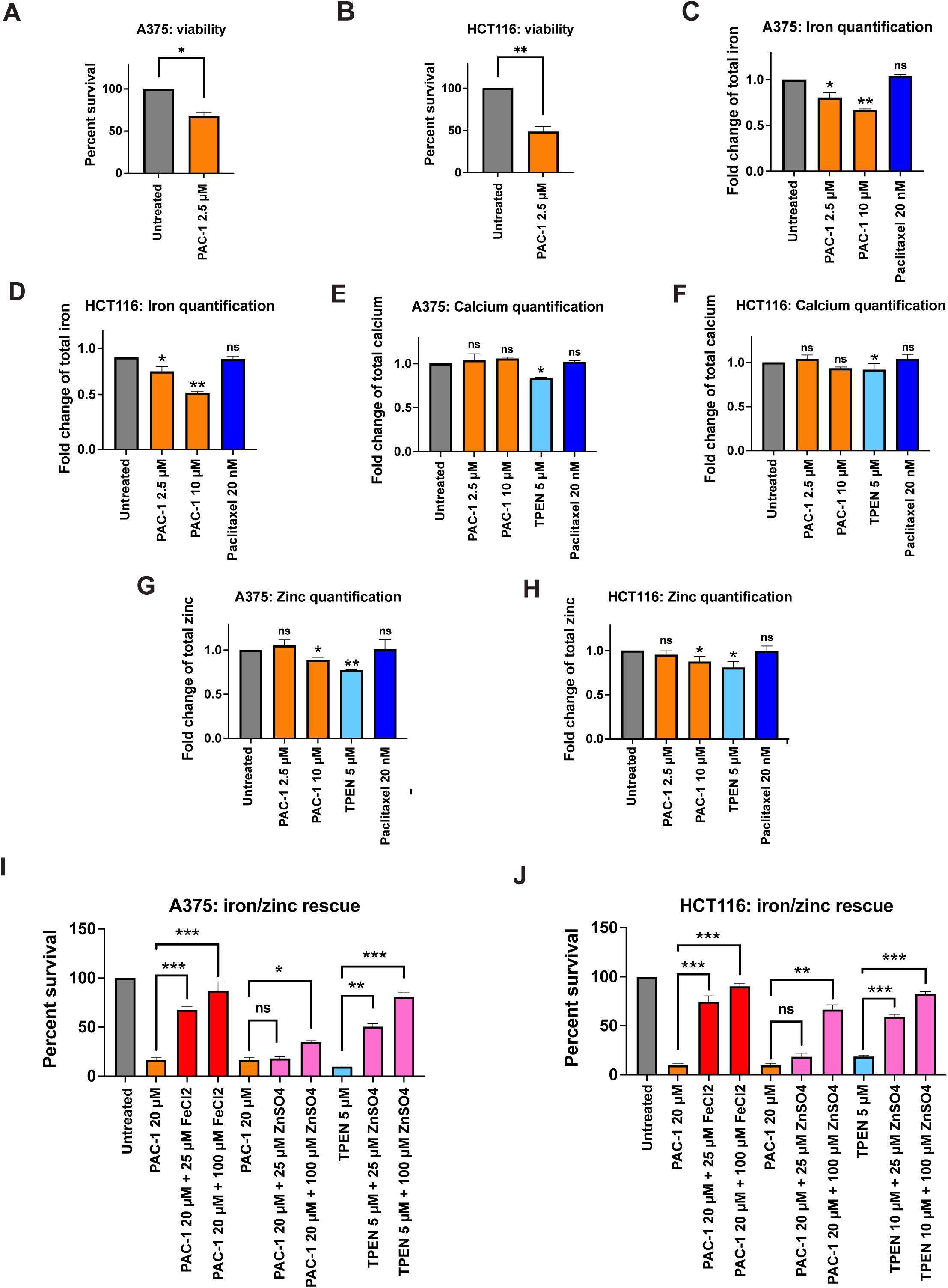
Cytotoxic concentrations of PAC-1 deplete iron but not other metals. (A) Quantification of cellular viability in A375 cells treated with 2.5 μM PAC-1. (B) Quantification of cellular viability in HCT116 cells treated with 2.5 μM PAC-1. (C) Quantification of cellular iron in A375 cells treated with 2.5 μM PAC-1, 10 μM PAC-1, or 20 nM paclitaxel. (D) Quantification of cellular iron in HCT116 cells treated with 2.5 μM PAC-1, 10 μM PAC-1, or 20 nM paclitaxel. (E) Quantification of cellular calcium in A375 cells treated with 2.5 μM PAC-1, 10 μM PAC-1, 5 μM TPEN, or 20 nM paclitaxel. (F) Quantification of cellular calcium in HCT116 cells treated with 2.5 μM PAC-1, 10 μM PAC-1, 5 μM TPEN, or 20 nM paclitaxel. (G) Quantification of cellular zinc in A375 cells treated with 2.5 μM PAC-1, 10 μM PAC-1, 5 μM TPEN, or 20 nM paclitaxel. (H) Quantification of cellular zinc in HCT116 cells treated with 2.5 μM PAC-1, 10 μM PAC-1, 5 μM TPEN, or 20 nM paclitaxel. (I) Quantification of cellular viability in A375 cells treated with 20 μM PAC-1 or 5 μM TPEN along with the indicated concentration of FeCl_2_ or ZnSO_4_. (J) Quantification of cellular viability in HCT116 cells treated with 20 μM PAC-1 or 5 μM TPEN along with the indicated concentration of FeCl_2_ or ZnSO_4_. *, p < .05; **, p < .005; ***, p < .0005; Student’s t-test (two-sided).

**Figure S4.**
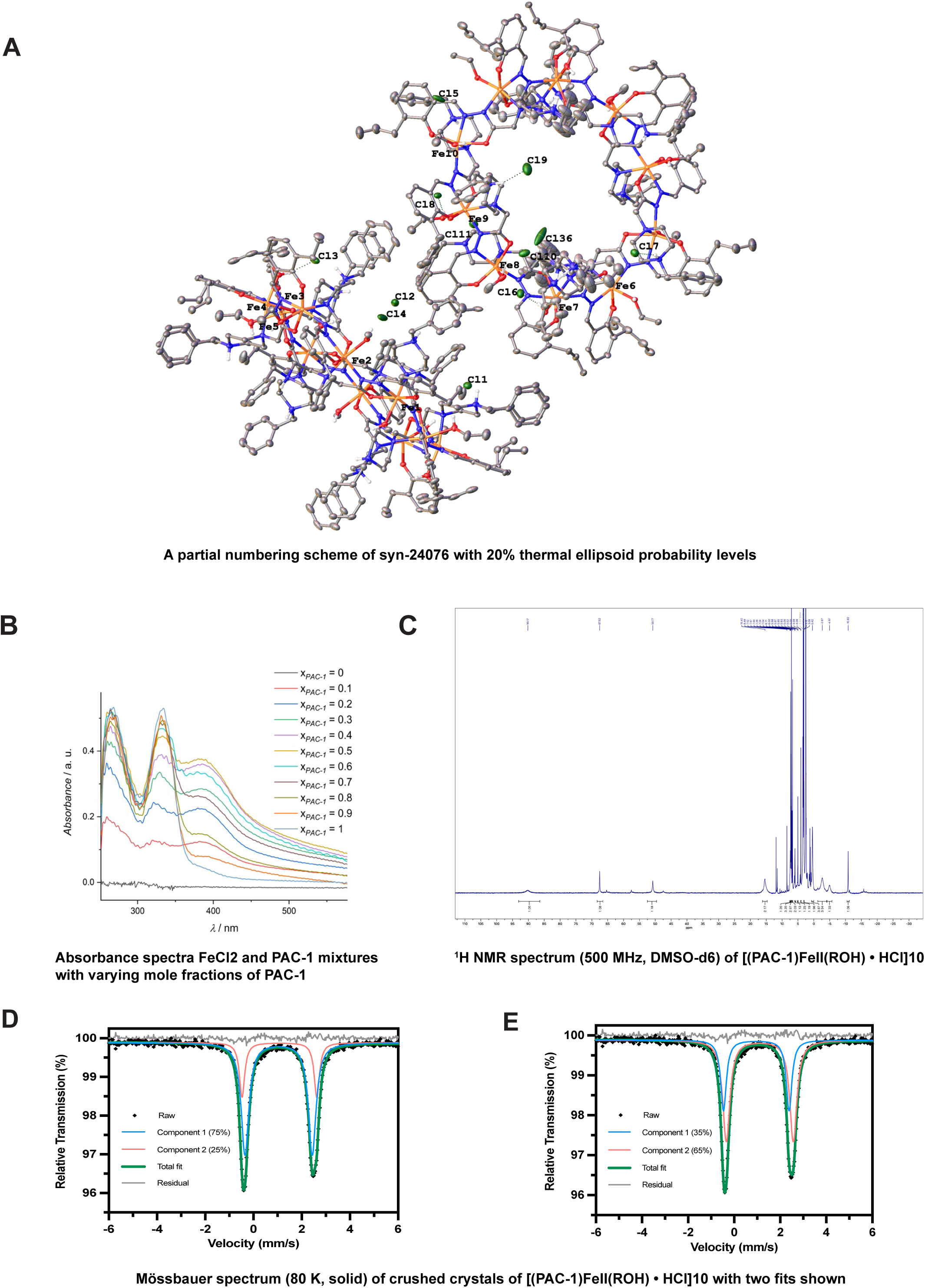
Structural analysis of the binding between PAC-1 and iron. (A) A partial numbering scheme of the PAC-1-iron complex (syn-24076) with 20% thermal ellipsoid probability levels. Carbon-bound hydrogen atoms are omitted for clarity. (B) Absorbance spectra of mixtures of FeCl_2_ and PAC-1 in buffer A (50 mM HEPES, 100 mM KNO3, pH = 7.4) with varying mole fractions of PAC-1. (C) ^1^H NMR spectrum (500 MHz, DMSO-d6) of [(PAC-1)FeII(ROH) • HCl]10. The large chemical shifts of the peaks are consistent with high-spin iron(II), in agreement with the Mössbauer spectrum below, but prevent assignment of resonances. In solution, the compound may break up into monomers with additional solvent molecules coordinated to the Fe centers. (D) Mössbauer spectrum (80 K, solid) of crushed crystals of [(PAC-1)FeII(ROH) • HCl]10. Raw data is shown in black diamonds, the component fits are shown in red and blue, the total fit is shown in bold green, and the residual is shown on top in gray. (E) Second fit for the Mössbauer spectrum (80 K, solid) of crushed crystals of [(PAC-1)FeII(ROH) • HCl]10. Raw data is shown in black diamonds, the component fits are shown in red and blue, the total fit is shown in bold green, and the residual is shown on top in gray.

**Figure S5.**
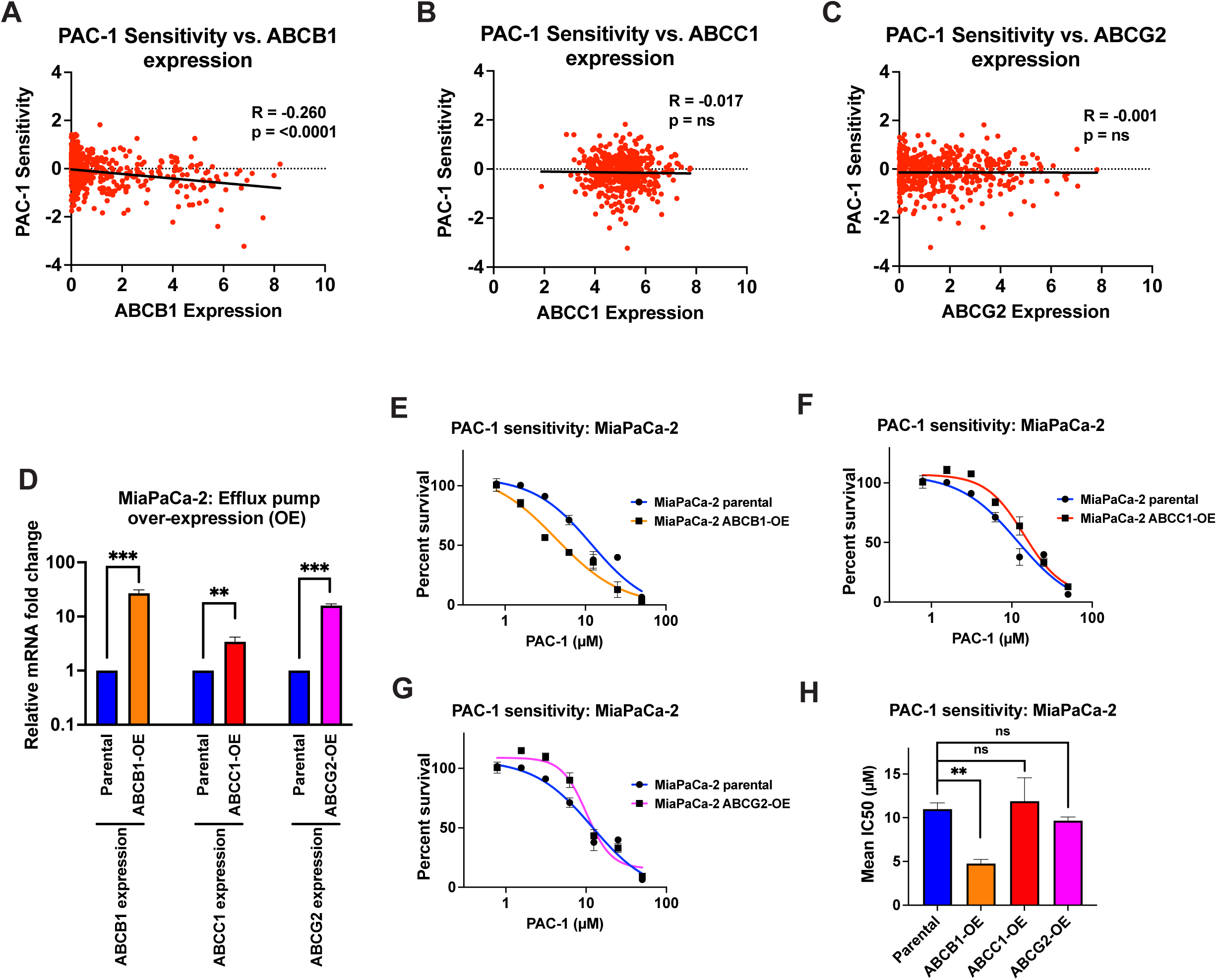
Overexpressing ABCC1 or ABCG2 does not affect PAC-1 sensitivity. (A) Scatter plot comparing the sensitivity of cancer cell lines to PAC-1 and the expression of ABCB1 (MDR1) in each cell line in the PRISM dataset. (B) Scatter plot comparing the sensitivity of cancer cell lines to PAC-1 and the expression of ABCC1 (MRP1) in each cell line in the PRISM dataset. (C) Scatter plot comparing the sensitivity of cancer cell lines to PAC-1 and the expression of ABCG2 (BCRP1) in each cell line in the PRISM dataset. (D) qPCR analysis of ABCB1, ABCC1, or ABCG2 expression in wild-type MiaPaCa-2 cells and in MiaPaCa-2 cells transduced with plasmids encoding each of these transporter genes. (E) 7-point dose response curves displaying cell viability in MiaPaCa-2 wild-type and ABCB1-overexpressing cells exposed to the indicated concentrations of PAC-1. (F) 7-point dose response curves displaying cell viability in MiaPaCa-2 wild-type and ABCC1-overexpressing cells exposed to the indicated concentrations of PAC-1. (G) 7-point dose response curves displaying cell viability in MiaPaCa-2 wild-type and ABCG2-overexpressing cells exposed to the indicated concentrations of PAC-1. (H) A bar graph summarizing the effects of transporter gene over-expression on PAC-1 sensitivity in MiaPaCa-2 cells. *, p < .05; **, p < .005; ***, p < .0005; Student’s t-test (two-sided).

**Figure S6.**
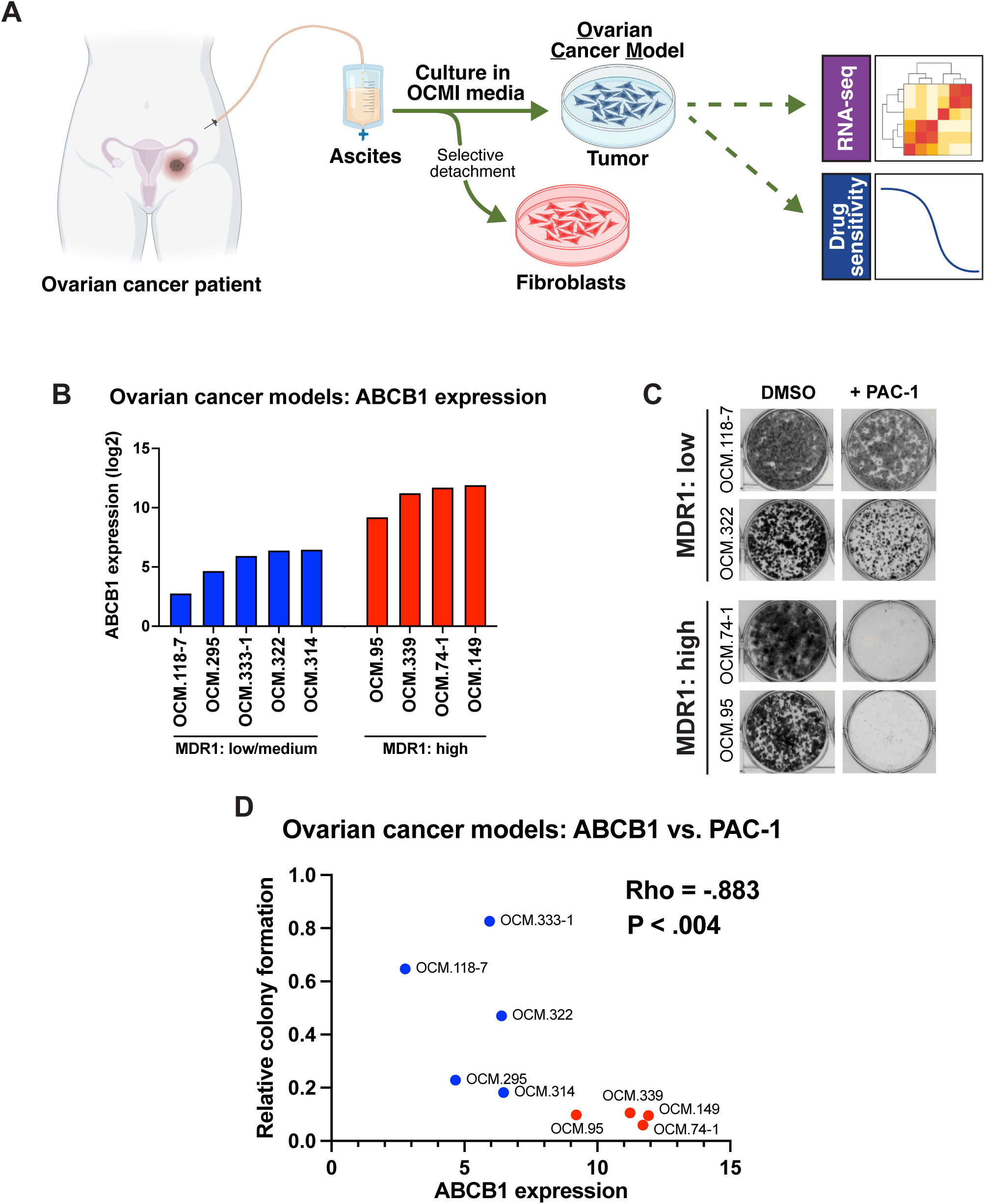
Testing the effects of PAC-1 in MDR1-high patient-derived ovarian cancer cells. (A) A schematic portraying how the patient-derived ovarian cancer cell models were obtained and characterized. (B) Quantification of ABCB1 expression in nine patient-derived OCMs by RNA sequencing. (C) Representative colony formation images following a 21-day exposure of the indicated OCMs to PAC-1. (D) A scatterplot displaying the relationship between ABCB1 expression and colony formation in nine patient-derived OCMs.

**Figure S7.**
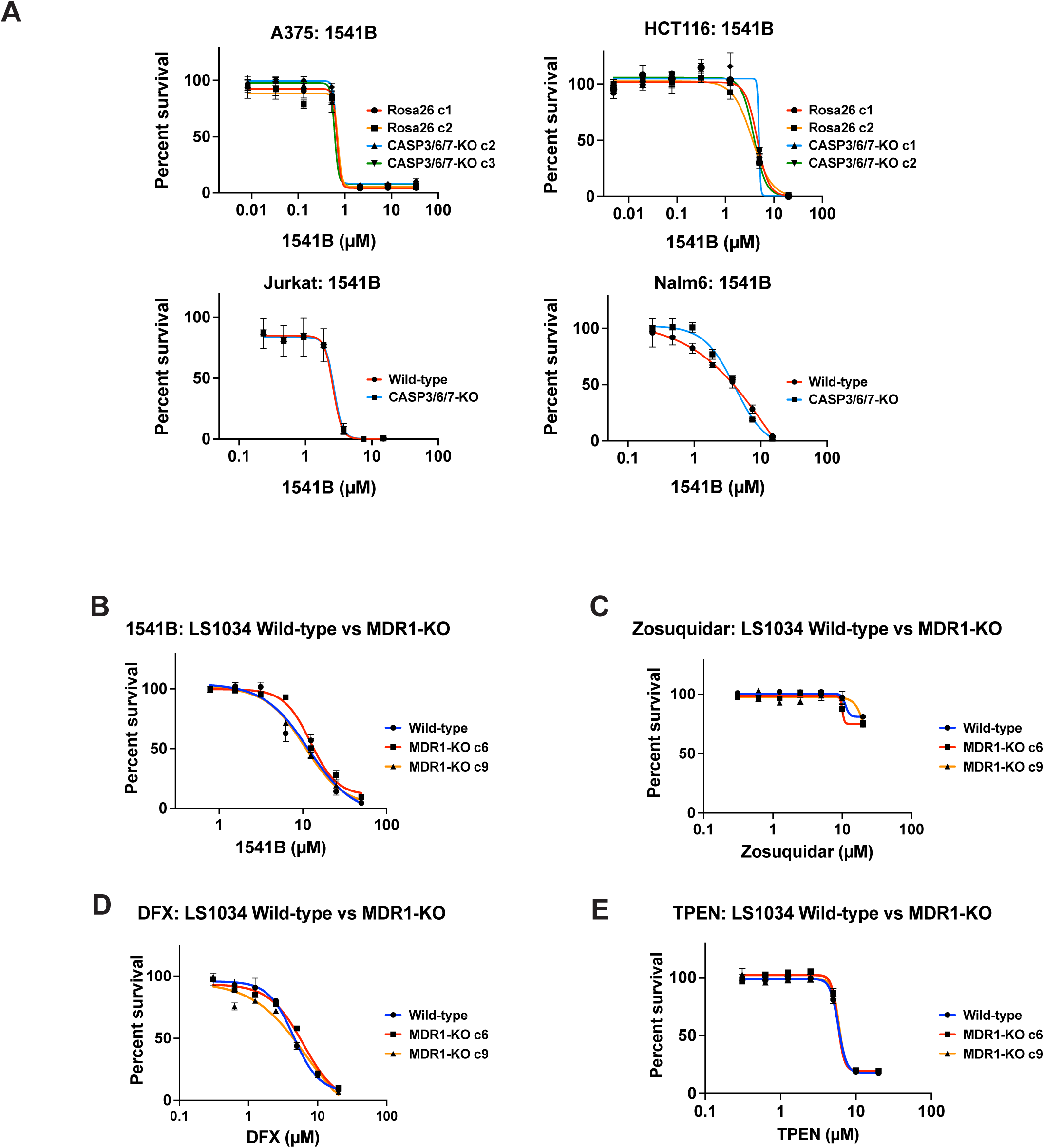
Other putative caspase-activating compounds and iron chelators do not exhibit MDR1 selectivity. (A) 7-point dose response curves displaying cell viability in control or CASP3/6/7-KO clones exposed to the indicated concentrations of 1541B. (B) 7-point dose response curves displaying cell viability in LS1034 wild-type and MDR1-KO clones exposed to the indicated concentrations of 1541B. (C) 7-point dose response curves displaying cell viability in LS1034 wild-type and MDR1-KO clones exposed to the indicated concentrations of zosuquidar. (D) 7-point dose response curves displaying cell viability in LS1034 wild-type and MDR1-KO clones exposed to the indicated concentrations of deferasirox (DFX). (E) 7-point dose response curves displaying cell viability in LS1034 wild-type and MDR1-KO clones exposed to the indicated concentrations of TPEN.

**Figure S8.**
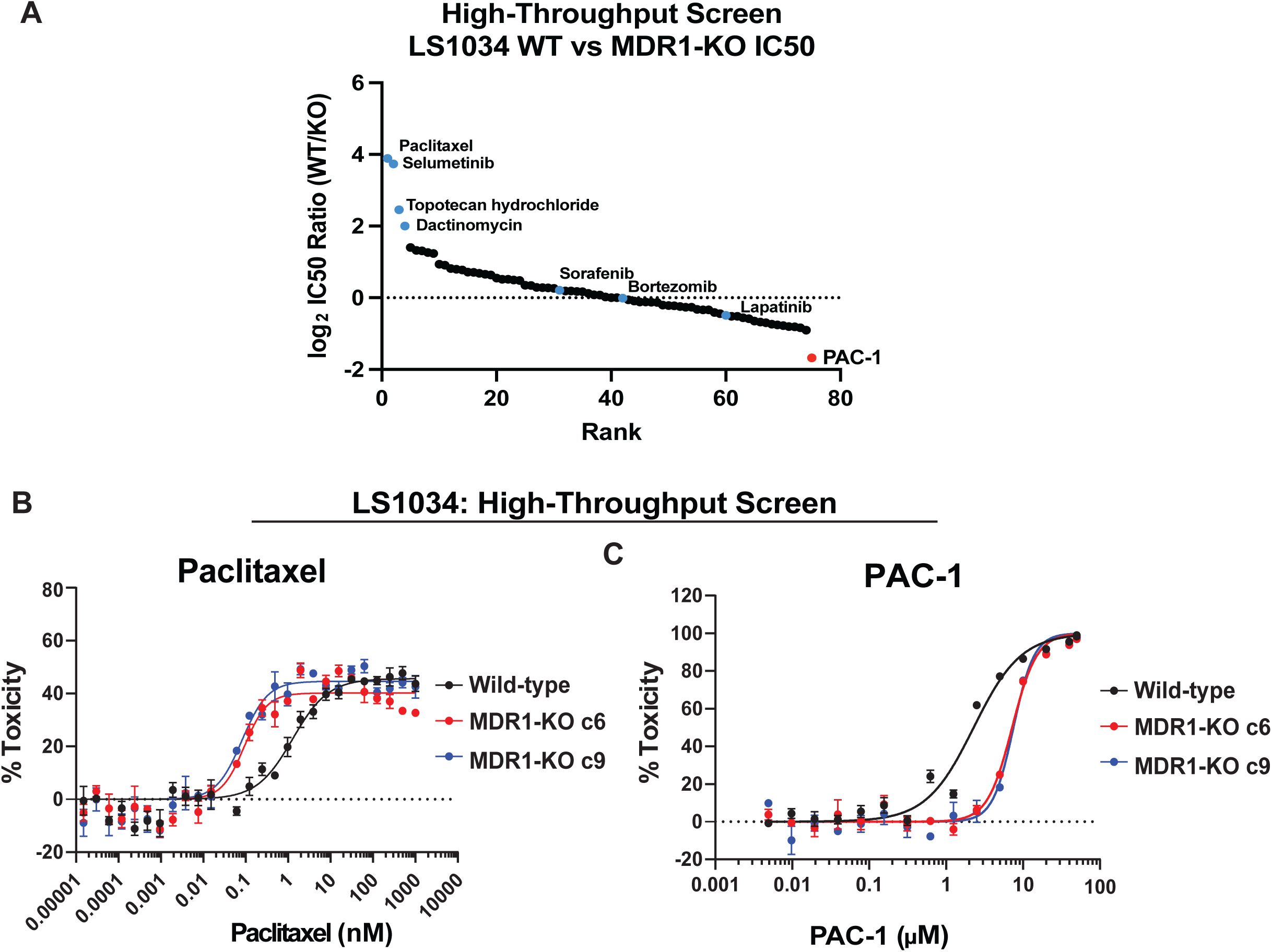
Screening approved and experimental cancer drugs for MDR1-selective cytotoxicity. (A) A waterfall plot displaying the ratio between the IC50 values of cancer drugs in LS1034 wild-type and MDR1-KO clones, arranged from highest to lowest. (B and C) Representative drug sensitivity curves generated in the high-throughput screen for paclitaxel and PAC-1.

## Notes

### Summary of Updates

Updated to include additional data and collaborators on MDR1 activity.

